# Paracrine Action of Bone Morphogenetic Protein 3 in Pulmonary Arterial Hypertension

**DOI:** 10.1101/2025.06.13.659638

**Authors:** Aymen Halouani, Eric Mensiah, Zaujia Athumani, Michael G. Katz, Katherine Jankowski, Maryam Mansoori, Vicki Rosen, Kiyotake Ishikawa, Lahouaria Hadri, Yassine Sassi

**Author notes:** Correspondence to: Yassine Sassi, PhD, Center for Vascular and Heart Research, Fralin Biomedical Research Institute at Virginia Tech Carilion, Roanoke, VA, 24016.

## Abstract

**Background:** Despite recent advancements in the management of pulmonary arterial hypertension (PAH), the disease remains devastating, with limited survival. Although the Bone Morphogenetic Protein (BMP) signaling pathway is known to play an important role in PAH, our current understanding of this pathway remains limited.

**Methods:** We assessed BMP3 levels in the lungs of mice, rats, and pigs with pulmonary hypertension, and in pulmonary vascular cells from human patients with PAH. We performed in vitro studies on human pulmonary artery smooth muscle cells (hPASMCs) and human pulmonary artery endothelial cells (hPAECs) derived from healthy donors and from patients with PAH. We generated mice with global or SMC-specific deletion of BMP3. Recombinant BMP3 protein and adeno-associated viruses (AAV) were used to overexpress BMP3 in two different models of PAH in rodents. Magnetic resonance imaging, cardiac hemodynamics, morphometric, and histological measurements were performed to evaluate the effects of BMP3 on cardiac function and pulmonary vascular remodeling.

**Results:** BMP3 is predominantly expressed in PASMCs and is downregulated in PAH. In vitro, conditioned medium from siRNA-BMP3-transfected hPASMCs increased hPAECs migration and proliferation, while PASMC-derived BMP3 inhibited PAH-diseased PAEC dysfunction. In both global and SMC-specific BMP3-deficient mice, exposure to a model of PAH exacerbated cardiac and pulmonary vascular remodeling in middle-aged mice. An intraperitoneally injected recombinant BMP3 prevented and reversed PAH in mice. A lung-targeted overexpression of BMP3, via AAV1-BMP3, reversed pulmonary vascular remodeling and inhibited cardiac dysfunction in mice and rats. Mechanistically, BMP3 activated the BMP/Smad1,5,8 pathway, inhibited the TGF-β/Smad2,3 pathway, and decreased the expression of cell cycle genes in hPAECs and in the lungs of BMP3-treated animals with PAH.

**Conclusions:** Our findings provide evidence that BMP3 overexpression attenuates pulmonary vascular remodeling and inhibits cardiac dysfunction by restoring the balance between the TGF-β and BMP pathways through a cell-cell communication mechanism, offering a novel therapeutic pathway for PAH.

**Clinical Perspective:** *What is New?:*

- BMP3 is downregulated in the lungs of mice, rats, and pigs with pulmonary hypertension and in pulmonary artery smooth muscle cells of patients with PAH.
- BMP3 acts as a paracrine factor between pulmonary vascular cells.
- Overexpression of BMP3 decreases pulmonary vascular remodeling and reverses cardiac dysfunction in PAH-diseased rodents.
- BMP3 acts by restoring the balance between the TGF-β/Smad2,3 pathway and the BMP/Smad1,5,8 pathway

*What Are the Clinical Implications?:*

- BMP3 represents a promising disease-modifying agent with potential for clinical translation in the treatment of PAH.
- Overexpressing BMP3 with Adeno-Associated Viruses has therapeutic potential to treat PAH.

## INTRODUCTION

Pulmonary arterial hypertension (PAH), a rare and progressive subtype of pulmonary hypertension (PH), leads to right heart failure and, ultimately, death.^1^ PH is defined by a mean pulmonary arterial pressure greater than 20 mmHg at rest, assessed via right heart catheterization.^2^ Despite recent advancements in PAH management, the disease remains devastating, with limited survival.^3,4^ The underlying mechanisms of PAH are still incompletely understood, but early features include pulmonary artery endothelial cell (PAEC) dysfunction and structural remodeling of the pulmonary vessels.^5,6^

The transforming growth factor (TGF)-β/bone morphogenetic protein (BMP) signaling pathway plays an important role in PAH. The TGF-β superfamily comprises two major protein classes: the BMP/GDF class and the TGF-β/activin/nodal class. These pathways engage both canonical (Smad1/5/8 and Smad2/3) and non-canonical signaling mechanisms, which vary depending on environmental conditions and cell types.^7^ Loss-of-function mutations in BMP receptor (BMPR)2 and its effectors underscore the central role of BMP signaling in the disease.^8,9^ Upon activation by BMP ligands, BMPR2 forms a heteromeric complex with BMPR1, which triggers the phosphorylation of Smad1, 5, and 8. This leads to the formation of a signaling complex with Smad4 that translocate to the nucleus, regulating the expression of target genes, including inhibitors of DNA-binding (Id) genes.^10^ In PAH, decreased BMPR2 expression and BMP signaling are observed in lung tissues, supporting the notion that BMP signaling serves a protective role.^11^ In contrast, TGF-β signaling, which involves Smad2 and Smad3 activation, is upregulated in human PAH and experimental PAH models. Inhibiting TGF-β signaling, such as through a selective TGF-β1/3 ligand trap, ameliorates experimental PH.^12^ Additionally, blocking activin signaling with Activin A Receptor Type 2 A (ACTRIIA)-Fc reduces pulmonary vascular remodeling.^13^ The balance between BMP and TGF-β signaling in pulmonary vascular cells is critical, and evidence suggests that PAH is characterized by a shift favoring TGF-β signaling.

While the BMP pathway is a promising target for the treatment of PAH, only a few BMP ligands have been studied in this fatal disease and few studies have been performed in animals. BMP4, a ligand secreted by endothelial cells in response to hypoxia, promotes PASMC proliferation and migration, contributing to pulmonary vascular remodeling.^14^ BMP2 is a homologous protein of BMP4, but it exerts a protective effect in PAH by inhibiting PASMC proliferation and migration.^15,16^ Opposing effects of BMP9 on pulmonary vascular remodeling and PAH have been reported.^17,18^ Although BMP3 (also known as Osteogenin) has been isolated from bone matrix, its expression is high in the developing lung and is, in adults, predominantly expressed in the lung.^19,20^ BMP3 is an important regulator of bone density and cancer progression.^21,22^ Nonetheless, the role of BMP3 in PAH remains to be investigated.

In this study, we explored the expression and effects BMP3 in PAH. We found that BMP3 is produced in PASMCs and is downregulated in the late phase of PAH. We show that PASMC-derived BMP3 inhibits PAEC proliferation and migration. SMC– specific Bmp3 deficiency resulted in a significant increase in right ventricular and pulmonary vascular remodeling in middle-aged mice. BMP3 upregulation, via the delivery of a recombinant protein or adeno-associated viruses (AAV), prevented and reversed PAH in mice and rats. This was mediated via the restoration of the balance between the TGF-β/Smad2,3 pathway and the BMP/Smad1,5,8 pathway. Our findings demonstrate an unprecedented role of BMP3 in PAH and underscore the therapeutic potential of BMP3 overexpression.

## METHODS

Source data for figures and supplemental figures are available from the corresponding author on reasonable request. Materials used to conduct the research are included in the article or the Methods in the Supplemental Material.

### Primary Human Pulmonary Vascular Cells

Primary human pulmonary artery smooth muscle cells (hPASMCs) and primary human pulmonary artery endothelial cells (hPAECs) isolated from healthy donors and from patients with idiopathic PAH were obtained from the Pulmonary Hypertension Breakthrough Initiative (PHBI). The patients’ characteristics are shown in the Supplemental Material (Tables S1 and S2). PASMCs and PAECs from healthy donors were also obtained from Lonza. All experiments were conducted using primary cells between passages two and seven. PASMCs were grown to 80% confluence before being treated with adenoviral vectors at a multiplicity of infection (MOI) of 30. The vectors included Ad-CMV-β-gal, Ad-BMP3, or Ad-shRNA-BMP3, all sourced from Vector Biolabs. Following a 48h incubation, the conditioned medium (CoM) was collected and transferred to PAECs in culture. Adenovirus-treated PASMCs were then harvested for RNA extraction to assess BMP3 expression level.

### Rat Hypoxia/Sugen PH model

Male Sprague-Dawley rats (8–10 weeks old) were used in this study. Pulmonary hypertension was induced by a single subcutaneous injection of SU5416 (20 mg/kg), followed by exposure to hypoxia (10% O₂) for three weeks. The normoxix control group was maintained under normoxic conditions (21% O₂). Three weeks after the initiation of hypoxia, rats exposed to sugen/hypoxia (Su/Hx) were randomly divided into two groups: AAV1-Luciferase (Su/Hx+AAV1-Ctrl) and AAV1-BMP3 (Su/Hx+AAV1-BMP3). The AAVs were delivered via IT instillation (3×10¹¹ CFU). Following the AAV treatment period, animals were maintained under normoxic conditions until the study endpoint.

### Statistical Analysis

All data are presented as mean ± standard error of the mean (SEM). Statistical analyses were performed using GraphPad Prism. Differences between two groups were analyzed using an unpaired two-tailed Student’s t-test, while comparisons among multiple groups were conducted using one-way or two-way ANOVA followed by Tukey’s post hoc test, as appropriate. Normality of data distribution was assessed using the Shapiro-Wilk test. A p-value < 0.05 was considered statistically significant. All experiments were performed with at least three independent biological replicates unless otherwise stated. For all in vivo experiments, a minimum of 5-8 mice per group was used to ensure statistical power. All experiments were performed with at least three independent biological replicates unless otherwise stated.

## RESULTS

### BMP3 is Downregulated in PAH

To determine whether BMP3 expression is altered in PAH, we quantified the expression levels of BMP3 in lung tissues from control and PAH-diseased mice exposed to chronic hypoxia (10% O_2_) for 3 weeks with a weekly injection of Sugen (a vascular endothelial growth factor receptor inhibitor). The analyses revealed a prominent decrease of pulmonary BMP3 mRNA levels in the lungs of PAH-diseased mice (Figure 1A). Next, we explored BMP3 expression profile in another *in vivo* PAH animal model: Sugen/Hypoxia-induced PAH in rats. Male rats received a single injection of Sugen and were then exposed to chronic hypoxia (10% O_2_) for 3 weeks. Rats exposed to normoxic conditions (21% O_2_) were used as controls. The PCR analyses revealed a significant decrease in BMP3 levels in diseased rats’ lungs (Figure 1B).

**Figure 1:**
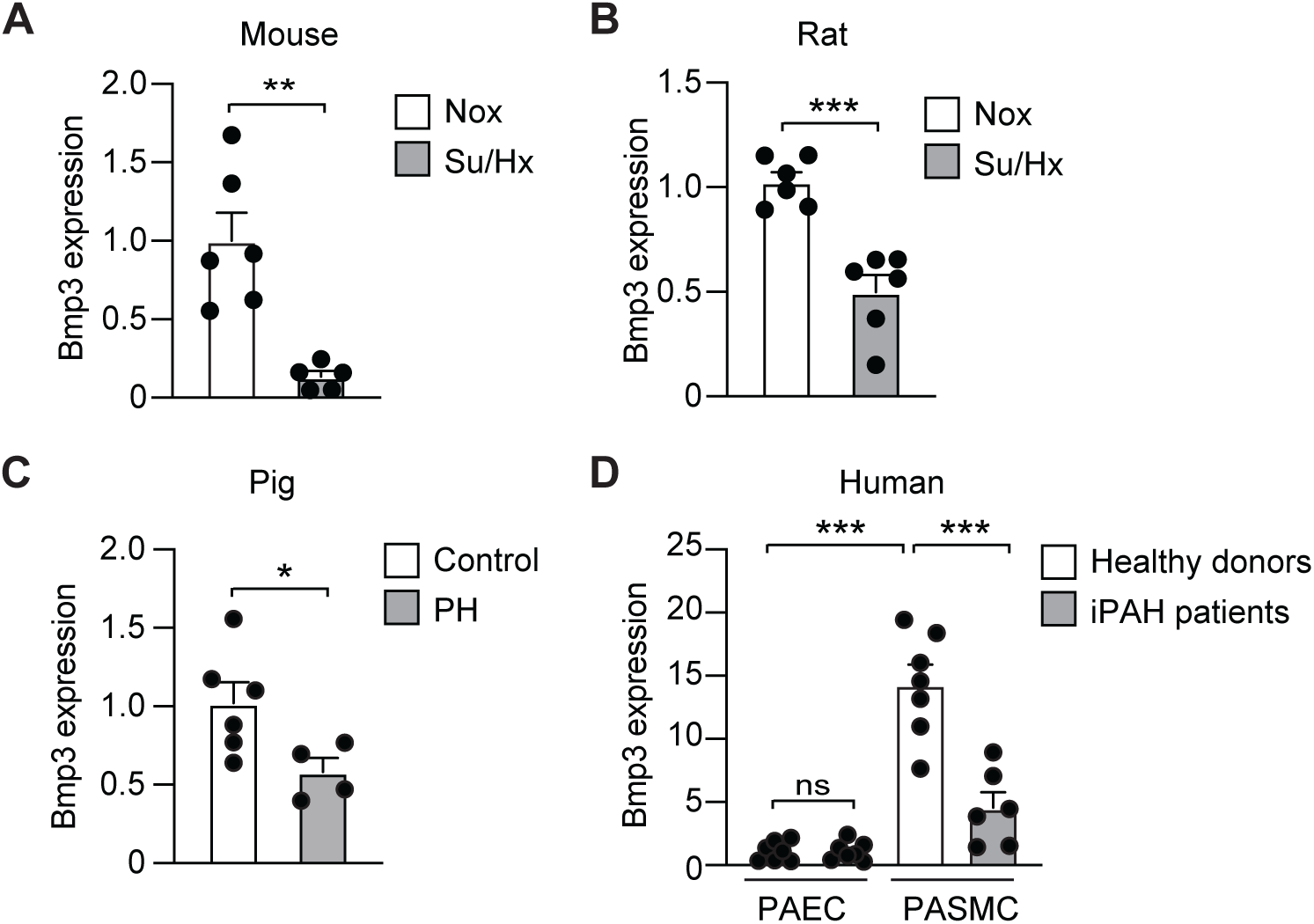
BMP3 regulation in PAH. **A**, Bmp3 mRNA expression level in lungs from healthy (normoxic) and PH (sugen/hypoxia) mice (n = 5-6 mice/group). **B,** Lung expression of Bmp3 in lung homogenates from normoxic and 3-week hypoxic male rats (n = 6 rats/group). **C,** Bmp3 expression in lungs from control and PV banding-induced PH pigs (n = 4-6 pigs/group). **D,** BMP3 mRNA levels, assessed by real-time PCR, in PAECs and PASMCs isolated from lungs of non-PAH and iPAH patients (n = 6-7 patients/group). Su/Hx: sugen/hypoxia. * P < 0.05, ** P < 0.01; *** P < 0.001, by two-tailed *t* test or 2-way ANOVA.

We next extended these observations to a large animal model using the swine model of post-capillary PH model.^26^ Small domestic pigs underwent thoracotomy with banding of the inferior pulmonary vein, which progressively increased pulmonary venous pressure and induced PH and right ventricular (RV) failure over four months (Table S3). Consistent with the rodent models, BMP3 levels were significantly reduced in the lungs of PH-diseased pigs (Figure 1C). We next quantified the expression levels of BMP3 in pulmonary artery endothelial cells (PAECs) and pulmonary artery smooth muscle cells (PASMCs) isolated from 7 non-PAH patients and a cohort of 7 patients with idiopathic PAH (iPAH, Tables S1 and S2). We found BMP3 levels, in cells isolated from healthy donors, to be substantially higher in PASMCs than in PAECs. Additionally, we found BMP3 expression to be significantly decreased in PASMCs isolated from the lungs of patients with clinical PAH (Figure 1D). These results indicate that BMP3 is predominantly expressed in PASMCs and is downregulated in PAH.

### PASMC-Derived BMP3 Inhibits PAEC Proliferation and Migration

We next investigated the effects of BMP3 on PASMC proliferation and migration. PASMCs (isolated from non-PAH donors) were treated with a recombinant BMP3 (rBMP3) protein or PBS (as a control). Serum stimulation increased the proliferation and migration of hPASMCs, and BMP3 treatment did not affect hPASMCs proliferation or migration, as assessed by BrdU proliferation and scratch wound assays (Figure 2A-B). In contrast, rBMP3 treatment significantly suppressed hPAEC proliferation and migration, indicating a cell-type-specific response (Figure 2C-D). These results suggest that BMP3 attenuates serum-induced hPAEC proliferation and migration.

**Figure 2:**
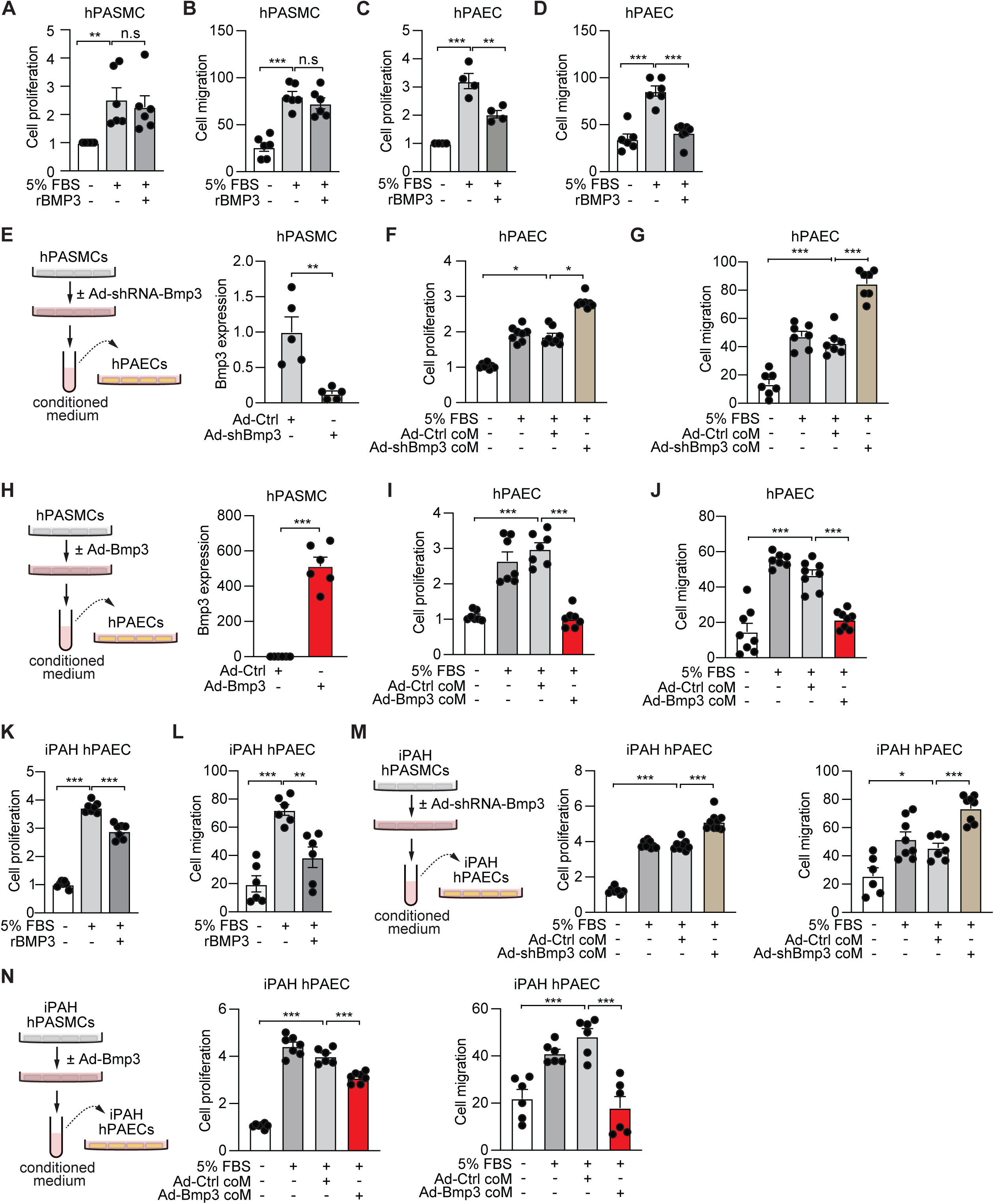
PASMC-derived BMP3 inhibits PAEC proliferation. **A**, Effect of BMP3 overexpression on hPASMC proliferation. Cells were treated with a recombinant BMP3 (rBMP3, 100 ng/ml) or PBS, and the proliferation was assessed 48 hours later. n = 6 experiments performed in triplicate. **B**, Effect of BMP3 upregulation on hPASMC migration. Cells were treated with a recombinant BMP3 (rBMP3, 100 ng/ml) or PBS and migration was assessed 48 hours later. n = 6 experiments performed in duplicate. **C**, Effect of BMP3 upregulation on hPAEC proliferation. Cells were treated with a recombinant BMP3 (rBMP3, 100 ng/ml) or PBS, and proliferation was assessed 48 hours later. n = 4 experiments performed in triplicate. **D**, Effect of BMP3 upregulation on hPAEC migration. Cells were treated with a recombinant BMP3 (rBMP3, 100 ng/ml) or PBS, and migration was assessed 48 hours later. n = 6 experiments performed in duplicate. **E**, (Left) Study design to assess cell-to-cell communication via BMP3. PASMCs were infected with Ad-shRNA-BMP3 or Ad-Ctrl. The conditioned medium (coM) from hPASMCs was then added to hPAECs, and the proliferation of hPAECs was assessed. (Right) BMP3 expression in hPASMCs treated with Ad-shRNA-BMP3 or Ad-Ctrl. n = 5 experiments performed in duplicate. **F**, Proliferation of hPAECs treated with Ad-shRNA-BMP3 coM or Ad-Ctrl coM. Proliferation was assessed 48 hours after the treatments. n = 8 experiments performed in triplicate. **G**, Migration of hPAECs treated with Ad-shRNA-BMP3 coM or Ad-Ctrl coM. n = 7 experiments performed in duplicate. **H**, (Left) Study design. PASMCs were infected with Adenovirus-GFP (Ad-Ctrl) or Adenovirus-BMP3 (Ad-BMP3; MOI 30 each). The conditioned medium from hPASMCs was then added to hPAECs, and the proliferation of hPAECs was assessed. (Right) Quantification of BMP3 expression in hPASMCs infected with Ad-Ctrl or Ad-BMP3. n = 6 experiments performed in duplicate. **I**, Proliferation of hPAECs treated with Ad-Ctrl coM or Ad-BMP3 coM. Proliferation was assessed 48h after the treatments. n = 7 experiments performed in triplicate. **J**, Migration of hPAECs treated with Ad-BMP3 coM or Ad-Ctrl coM. n = 8 experiments performed in duplicate. **K-L**, Effect of BMP3 upregulation on the proliferation (**K**) and migration (**L**) of hPAEC isolated from patients with idiopathic PAH (iPAH). n = 6-7 experiments performed in duplicate. **M**, (Left) Study design to assess cell-to-cell communication in iPAH pulmonary vascular cells via BMP3. iPAH-PASMCs were infected with Ad-shRNA-BMP3 or Ad-Ctrl. The conditioned medium (coM) was then added into iPAH-PAECs, and the proliferation and migration of iPAH-PAECs were assessed. (Middle) Proliferation and (Right) migration of iPAH-PAECs treated with Ad-shRNA-BMP3 coM or Ad-Ctrl coM. n = 6-9 experiments. **N**, (Left) Study design. iPAH-PASMCs were infected with Adenovirus-GFP (Ad-Ctrl) or Adenovirus-BMP3 (Ad-BMP3; MOI 30 each). The conditioned medium was then added into iPAH-PAECs, and the proliferation and migration of iPAH-hPAECs were assessed. (Middle) Proliferation and (Right) migration of iPAH-PAECs treated with Ad-Ctrl coM or Ad-BMP3 coM. n = 6-7 experiments performed in triplicate. * P < 0.05; ** P < 0.01; *** P < 0.001. ns: not significant, by two-tailed *t* test or one-way ANOVA. FBS: Fetal Bovine Serum.

Given that BMP3 is predominantly expressed in PASMCs, we next investigated whether BMP3-derived hPASMC acts in a paracrine manner to influence PAEC function. In a first approach, we assessed the effects of endogenous BMP3 using a specific siRNA designed against human BMP3. Two days after siRNAs transfection in hPASMCs, the medium was collected and transferred into hPAECs (Figure 2E). Si-BMP3 transfection in hPASMCs led to a 70% reduction in BMP3 mRNA level, showing the efficiency of siRNA-BMP3 (Figure 2E). Conditioned medium from BMP3-silenced PASMCs significantly increased hPAEC proliferation and migration compared to conditioned medium from siRNA control-transfected PASMCs (Figure 2F-G), suggesting that endogenous PASMC derived-BMP3 inhibits PAEC activation.

To complement this approach, we next tested the effect of conditioned medium from hPASMCs that overexpress BMP3 on hPAECs proliferation and migration. PASMCs were treated with an adenovirus encoding GFP (Ad-Ctrl) or BMP3 (Ad-Bmp3) in the presence of 5% Serum. Two days later, the conditioned medium was collected and transferred into hPAECs (Figure 2H). Compared to the control Ad-GFP, increased BMP3 levels (Figure 2H), significantly reduced hPAEC proliferation and migration (Figure 2I-J), further confirming BMP3 paracrine inhibitory effects.

We next used hPASMCs and hPAECs derived from patients with iPAH. Recombinant BMP3 treatment induced a significant reduction in PAH-PAECs proliferation and migration (Figure 2 K-L). Conditioned medium from BMP3-silenced PAH-PASMCs enhanced PAH-PAEC proliferation and migration (Figure 2M), while BMP3 overexpression in PAH-PASMCs led to the opposite effect, significantly inhibiting PAH-PAEC proliferation and migration (Figure 2N). Together, these results indicate that BMP3 is a key paracrine mediator involved in cell-to-cell communication between PASMC and PAEC, where BMP3-derived PASMC inhibits PAEC proliferation and migration. This intracellular signaling communication mechanism mediated by BMP3 is impaired in PAH and may contribute to pathological vascular remodeling.

### BMP3 Deletion Exacerbates PAH in Middle-Aged Mice

To investigate the in vivo role of BMP3 in PH, we generated global Bmp3 KO mice by crossing conditional Bmp3^flox/flox^ mice with UBC-CreER^T2^ mice and subjected them to the Su/Hx-induced PH model. In young (3-month-old) mice, global *BMP3* deletion did not alter the PH phenotype. Right ventricular systolic pressure (RVSP) and pulmonary vascular remodeling were comparable between BMP3 KO and WT mice under normoxic and hypoxic conditions (Figure 3A-E). Both young male and female global Bmp3 KO mice displayed the same phenotype outcome (Figure S1).

**Figure 3:**
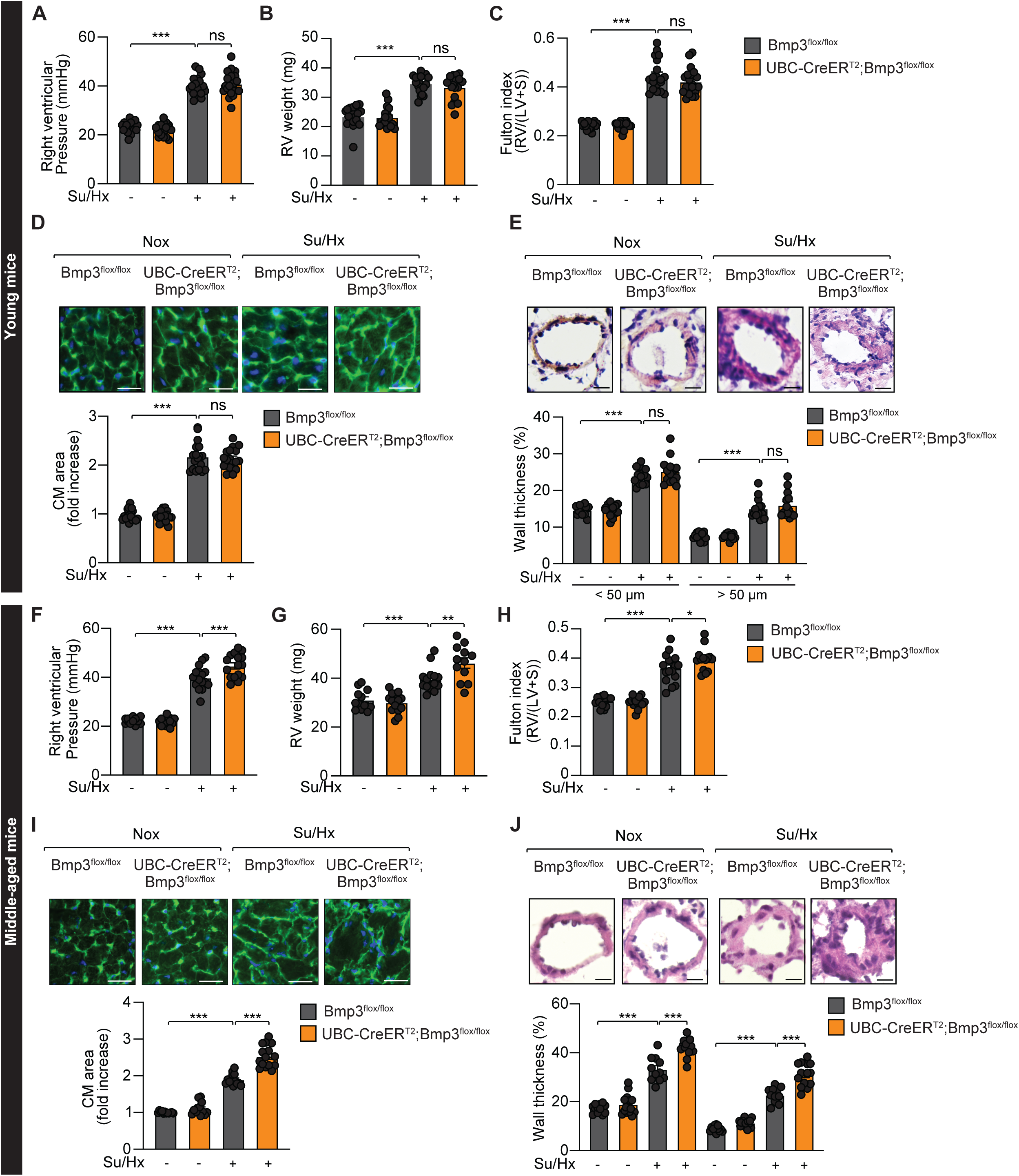
Global BMP3 deletion exacerbates PH in middle-aged mice. Bmp3^flox/flox^ mice were crossed with UBC-CreER^T2^ mice. The offspring mice were then injected with tamoxifen at 6-8 weeks or 9-12 months of age for 5 consecutive days. Two weeks later, mice received 20 mg/kg of SU5416 (SU), and were then exposed to 4 weeks of hypoxia (Hx). SU5416 was injected once a week during the next 2 weeks. The endpoint for hemodynamic measurements and sacrifice was 4 weeks after the first Sugen injection. **A-E**, Hemodynamic, morphometric and histological measurements in young mice. **A**, Right ventricular systolic pressure (RVSP) of the indicated groups. n = 18-21 mice/group. **B**, Right Ventricular (RV) weight of the indicated groups. n = 18-21 mice/group. **C**, Fulton index of the indicated groups. n = 18-21 mice/group. **D**, (Upper panel) Representative WGA-stained RV sections to assess hypertrophy of cardiac myocytes. Scale bar: 50 μm. (Lower panel) Quantitative analysis. n = 18-21 mice/group. **E**, (Upper panel) Representative H&E-stained pulmonary artery sections from the indicated groups. (Lower panel) Percentage of arterial wall thickness. **F-J**, Hemodynamic, morphometric and histological measurements in middle-aged mice. **F**, RVSP of the indicated groups. n = 14-17 mice/group. **G**, RV weight of the indicated groups. n = 14-17 mice/group. **H**, Fulton index of the indicated groups. n = 14-17 mice/group. **I**, (Upper panel) Representative WGA-stained RV sections to assess hypertrophy of cardiac myocytes. Scale bar: 50 μm. (Lower panel) Quantitative analysis. n = 14-17 mice/group. (Upper panel) **J,** Representative H&E-stained pulmonary artery sections from the indicated groups. (Lower panel) Percentage of arterial wall thickness. n = 14-17 mice/group. * P < 0.05; ** P < 0.01; *** P < 0.001; ns: not significant by 2-way ANOVA.

In middle-aged (9–12-month-old) mice, global Bmp3 deletion under normoxic conditions also did not affect RV and pulmonary vascular remodeling compared to age-matched WT littermate controls (Figure 3F-J). However, Hypoxia (Su/Hx) exposure, middle-aged BPM3-deficient mice developed significantly more severe PH characterized by increased RVSP and RV hypertrophy compared to WT mice (Figure 3F-H), enhanced cardiomyocyte hypertrophy (Figure 3I), and pulmonary vascular remodeling (Figure 3J) compared littermate WT. These results demonstrated age-dependent effects of BMP3 by showing that a global deletion of BMP3 sensitizes and exacerbates PH in middle-aged mice, but not in young animals.

To determine whether the observed phenotype is attributable to BMP3 loss in SMC, we crossed Bmp3^flox/flox^ mice with SM22-Cre mice to generate SMC-specific BMP3 KO mice. Young mice with SMC-specific Bmp3 deletion displayed cardiac and pulmonary vascular remodeling comparable to that observed in their control littermates (Figure 3A-E). Young male and female SMC-specific deficient Bmp3 mice displayed the same phenotype (Figure S2). However, the middle-aged SMC-specific Bmp3 KO mice exhibited an exacerbated PH phenotype with elevated RVSP and RV hypertrophy (both at the tissue and cellular levels), and increased pulmonary vascular remodeling compared to control littermates (Figure 4F-J). The PH phenotype in middle-aged SMC-specific Bmp3 mice was similar to that of global Bmp3 KO mice. No sex-specific differences were observed in SMC-specific Bmp3 KO mice (Figure S3). Collectively, these findings demonstrate that BMP3-derived SMC plays a protective role in maintaining pulmonary vascular homeostasis and function in aging, and its loss exacerbates PH under pathological stress conditions.

**Figure 4:**
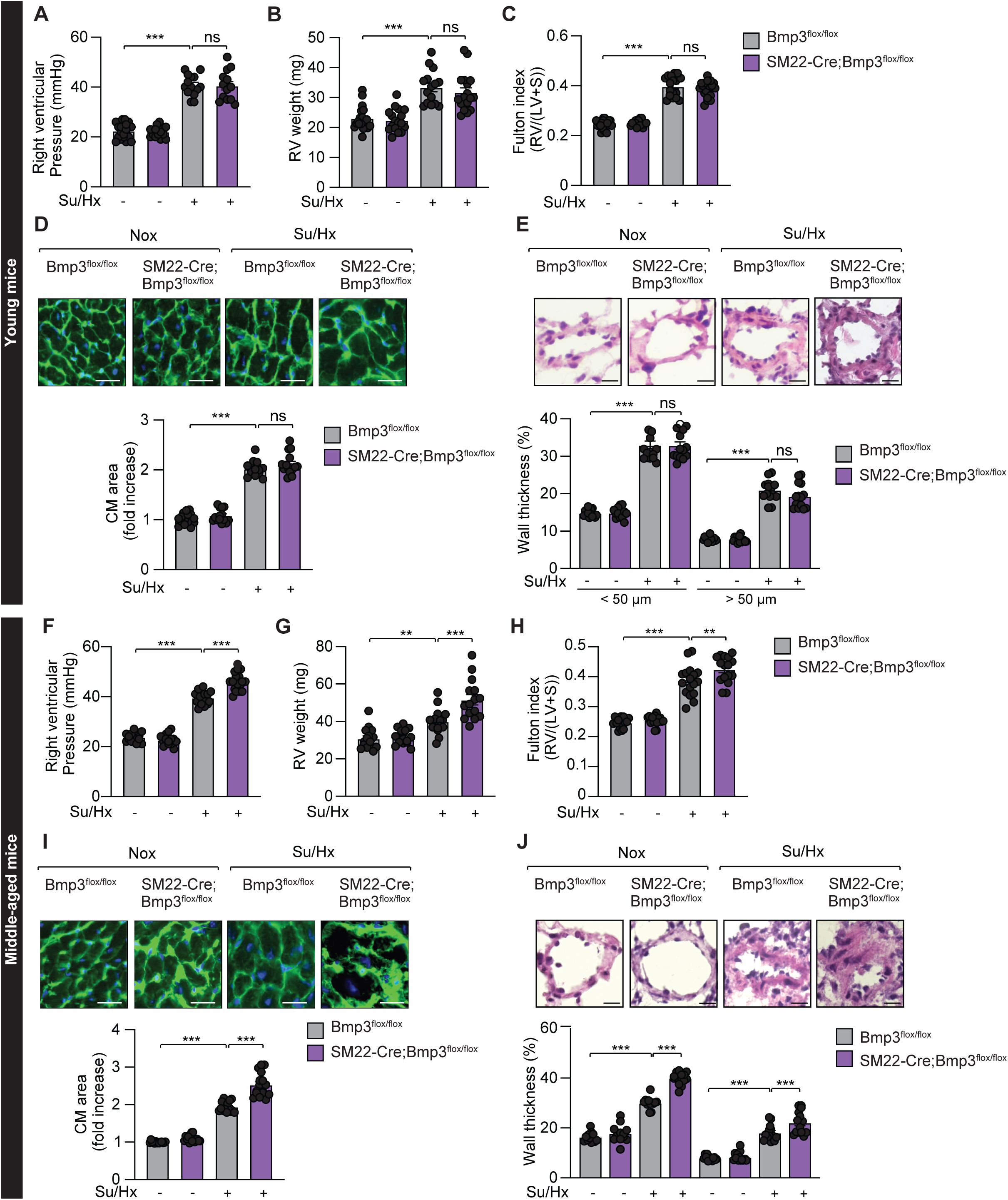
A Smooth-muscle cell-specific BMP3 deletion exacerbates PH in middle-aged mice. Bmp3^flox/flox^ mice were crossed with SM22-Cre mice. The offspring mice received 20 mg/kg of SU5416 (SU), and were then exposed to 4 weeks of hypoxia (Hx). SU5416 was injected once a week during the next 2 weeks. The endpoint for hemodynamic measurements and sacrifice was 4 weeks after the first Sugen injection. **A-E**, Hemodynamic, morphometric and histological measurements in young mice. **A**, Right ventricular systolic pressure (RVSP) of the indicated groups. n = 17-18 mice/group. **B**, Right Ventricular (RV) weight of the indicated groups. n = 17-18 mice/group. **C**, Fulton index of the indicated groups. n = 17-18 mice/group. **D**, (Upper panel) Representative WGA-stained RV sections to assess hypertrophy of cardiac myocytes. Scale bar: 50 μm. (Lower panel) Quantitative analysis. n = 17-18 mice/group. **E**, (Upper panel) Representative H&E-stained pulmonary artery sections from the indicated groups. (Lower panel) Percentage of arterial wall thickness. n = 17-18 mice/group. **F-J**, Hemodynamic, morphometric and histological measurements in middle-aged mice. **F**, RVSP of the indicated groups. n = 15-17 mice/group. **G**, RV weight of the indicated groups. n = 15-17 mice/group. **H**, Fulton index of the indicated groups. n = 15-17 mice/group. **I**, (Upper panel) Representative WGA-stained RV sections to assess hypertrophy of cardiac myocytes. Scale bar: 50 μm. (Lower panel) Quantitative analysis. n = 15-17 mice/group. **J**, (Upper panel) Representative H&E-stained pulmonary artery sections from the indicated groups. (Lower panel) Percentage of arterial wall thickness. n = 15-17 mice/group. * P < 0.05; ** P < 0.01; *** P < 0.001; ns: not significant by 2-way ANOVA.

### Exogenous Recombinant BMP3 Prevents and Inhibits Sugen/Hypoxia-induced PAH in Mice

To explore the therapeutic potential of BMP3 in PAH, we evaluated whether recombinant BMP3 (rBMP3) administration could prevent or reverse PAH induced by Su/Hx-exposure in mice for 3 weeks. In a preventive approach, WT C57B6 male mice were exposed to sugen and chronic hypoxia, and were IP administered rBMP3 (0.3 mg/kg) or vehicle saline every 3 days (Figure S4A). Saline-treated mice developed hallmark features of PAH, including elevated RVSP, RV weight, and Fulton index (Figure S4B-D). In contrast, mice treated with rBMP3 exhibited a marked reduction in all of these parameters, indicating a protective effect of rBMP3 against PAH development (Figure S4B-D). Furthermore, histological analyses showed that rBMP3 treatment prevented RV cardiomyocyte (CM) hypertrophy (Figure S4E) and significantly reduced the percentage of pulmonary arterial wall thickness (Figure S4F). These results indicate that rBMP3 prevents both RV and pulmonary vascular remodeling associated with PAH in the Su/Hx mouse model.

Next, a separate cohort of WT mice were exposed to Su/Hx for 3 weeks to establish PAH, and were then randomized to receive rBMP3 (0.3mg/kg, every 3 days) or saline for an additional 3 weeks (Figure S4G). rBMP3 treatment significantly decreased RVSP and RV hypertrophy compared to saline-treated controls (Figure S4H-J). At the cellular level, rBmp3 attenuated cardiomyocyte hypertrophy (Figure S4K). In addition, morphometric analysis demonstrated a marked decrease in pulmonary arterial wall thickness in rBMP3-treated mice (Figure S4L). These results demonstrate that a systemic administration of exogenous rBMP3 has both preventive and therapeutic effects in experimental Su/Hx-induced PAH in mice.

### Lung-Targeted BMP3 Gene Therapy Prevents and Reverses PAH in Mice

To evaluate the therapeutic efficacy of lung-targeted Bmp3 gene delivery *in vivo*, we employed an intratracheal delivery approach of adeno-associated virus serotype 1 (AAV1) (Figure 5A). AAV1-Bmp3 was successfully generated to achieve sustained and long-term expression of BMP3 in the lungs (Figure 5B). Next, randomized WT male mice were intratracheally instilled with AAV1-Bmp3 or AAV1 encoding luciferase as a control (2×10^11^ genome copies per mouse) two weeks prior to 4 weeks of exposure to hypoxia and weekly Sugen injections (Figure 5A). Consistent with our findings in recombinant Bmp3-treated mice, AAV1-Bmp3 treatment significantly reduced RVSP and RV hypertrophy induced by Sugen/hypoxia (Figure 5C-D). AAV1-Bmp3 attenuated RV cardiomyocyte hypertrophy and pulmonary arterial wall thickness (Figure 5E-F), confirming the broad preventive effects of BMP3 gene delivery. In accordance with the results obtained in male mice, AAV1-Bmp3 attenuated right ventricular dysfunction and pulmonary vascular remodeling in female mice (Figure S5). Together, these results indicate that a lung-targeted Bmp3 overexpression using intratracheal gene delivery of AAV1-Bmp3 prevents Sugen/Hypoxia-induced PAH in mice.

**Figure 5:**
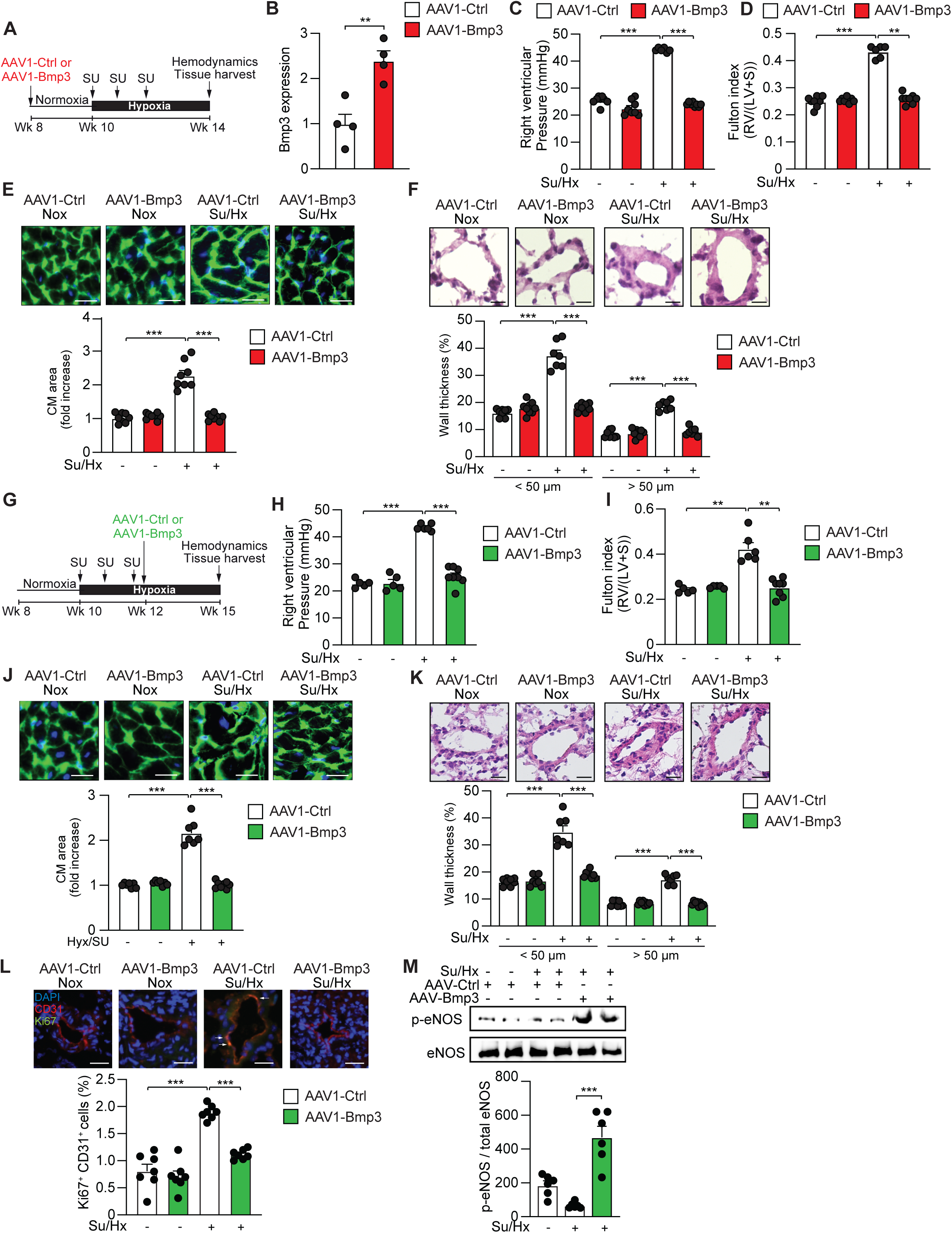
A lung-specific overexpression of BMP3 prevents and inhibits PH in mice. **A,** Design of the study. Male mice were randomly assigned to receive AAV1-Ctrl or AAV1-Bmp3 at week 8. Two weeks later, mice received 20 mg/kg of SU5416, and were then exposed to four weeks of chronic hypoxia. SU5416 was injected once a week during the next two weeks. The endpoint for hemodynamic measurements and sacrifice was at week 14. **B,** Bmp3 mRNA expression in lung homogenates from the indicated groups. n = 4 mice/group. **C**, RVSP of the indicated groups. n = 6-8 mice/group. **D**, Fulton index of the indicated groups. n = 6-8 mice/group. **E**, (Upper panel) Representative WGA-staining of RV sections. Scale bar: 50 μm. (Lower panel) Quantitative analysis. n = 6-8 mice/group. **F**, (Upper panel) Representative H&E-stained pulmonary artery sections from the indicated groups. Scale bar: 50 μm. (Lower panel) Percentage of arterial wall thickness in relation to cross-sectional diameter. n = 6-8 mice/group. **G,** Design of the study. Male mice received 20 mg/kg of SU5416 and were then exposed to two weeks of chronic hypoxia. SU5416 was injected once a week during the next two weeks. Mice were then randomly assigned to receive AAV1-Ctrl or AAV1-Bmp3 at week 12. The endpoint for hemodynamic measurements and sacrifice was at week 16. **H**, RVSP of the indicated groups. n= 5-8 mice/group. **I**, Fulton index of the indicated groups. n= 5-8 mice/group. **J**, (Upper panel) Representative WGA-staining of RV sections. Scale bar: 50 μm. (Lower panel) Quantitative analysis. n= 5-8 mice/group. **K**, (Upper panel) Representative H&E-stained pulmonary artery sections from the indicated groups. Scale bar: 40 μm. (Lower panel) Percentage of arterial wall thickness in relation to cross-sectional diameter. n= 5-8 mice/group. **L**, Ki-67/CD31 positive cells in normoxic and Su/Hx lungs treated with AAV1-Ctrl or AAV1-Bmp3. n= 7 mice/group. Scale bar: 25 μm **M**, Phosphorylated eNOS and total eNOS protein expression in normoxic and Su/Hx lungs treated with AAV1-Ctrl or AAV1-Bmp3. n= 6 mice/group. ** P < 0.01; *** P < 0.001, by two-tailed t test, 1-way ANOVA, or 2-way ANOVA. RV: Right ventricle, Su/Hx: Sugen/Hypoxia, CM: Cardiomyocyte.

To evaluate the therapeutic potential of AAV1-Bmp3 gene delivery *in vivo*, male mice were first exposed to Su/Hx conditions for 2 weeks, after which they were randomized to receive an intratracheal delivery of AAV1-BMP3 or AAV1-Ctrl treatment for 4 weeks (Figure 5G). Remarkably, AAV1-Bmp3-treated animals were protected from Su/Hx-induced PAH features, as they showed a marked decrease in RVSP and RV hypertrophy (Figure 5H-I), a reduction of cardiomyocyte hypertrophy (Figure 5J), and pulmonary vascular remodeling (Figure 5K). These effects were reproduced in female mice, confirming the efficacy of AAV1-BMP3 treatment across sexes (Figure S6). Moreover, immunofluorescence staining revealed that PAH-induced endothelial cell proliferation, as shown by increased CD31+ (EC-specific marker) and Ki67+ (a marker of cell proliferation) cells, was significantly reduced by BMP3 overexpression (Figure 5L). Furthermore, the beneficial effects of Bmp3 gene transfer were associated with restored eNOS phosphorylation at Ser1177, a key indicator of endothelial function, which was otherwise suppressed in PH (Figure 5M). These results indicate that a lung-targeted inhaled AAV1-BMP3 inhibits Sugen/Hypoxia-induced PAH in mice.

### BMP3 Overexpression Preserves the Cardiac Function and Inhibits Pulmonary Vascular Remodeling in a Severe Rat Model of PAH

We then determined whether BMP3 overexpression could reverse cardiac and pulmonary vascular remodeling in a severe model of PAH. We used the Su/Hx rat model because it exhibits more severe cardiac and pulmonary vascular remodeling than the Su/Hx model in mice.^27^ Rats received a single injection of Sugen followed by 3 weeks of chronic hypoxic conditions (10% O_2_) (Figure 6A). At day 21, rats were randomized to receive AAV1-Bmp3 or AAV1-Ctrl (3×10^11^ genome copies per rat) by intratracheal instillation, then returned to normoxic conditions for 4 weeks (Figure 6A). As expected, Su/Hx treatment induced a decrease in pulmonary Bmp3 level, which was restored by AAV1-Bmp3 lung delivery in PH-diseased rats (Figure 6B). Serial functional measures of cardiac function by magnetic resonance imaging revealed improved RV function with higher tricuspid annular plane systolic excursion (TAPSE), reduced RV chamber width, increased RV ejection fraction and higher cardiac index in AAV1-Bmp3-treated rats as compared with AAV1-control-treated rats (Figure 6C). Moreover, these improvements were accompanied by reduced pulmonary vascular resistance (PVR) and mean pulmonary artery pressure (mPAP) in AAV1-Bmp3-treated rats (Figure 6C), indicating improved cardiopulmonary function in rats upon AAV1-Bmp3 treatment. Together, the MRI analyses indicated improved cardiac function and decreased pulmonary vascular remodeling in rats upon AAV1-Bmp3 treatment. Invasive hemodynamic measurements demonstrated that AAV1-Bmp3-treated rats exhibited significantly reduced RVSP and mean pulmonary arterial pressures, when compared to AAV1-control-treated rats (Figure 6D-F).

**Figure 6:**
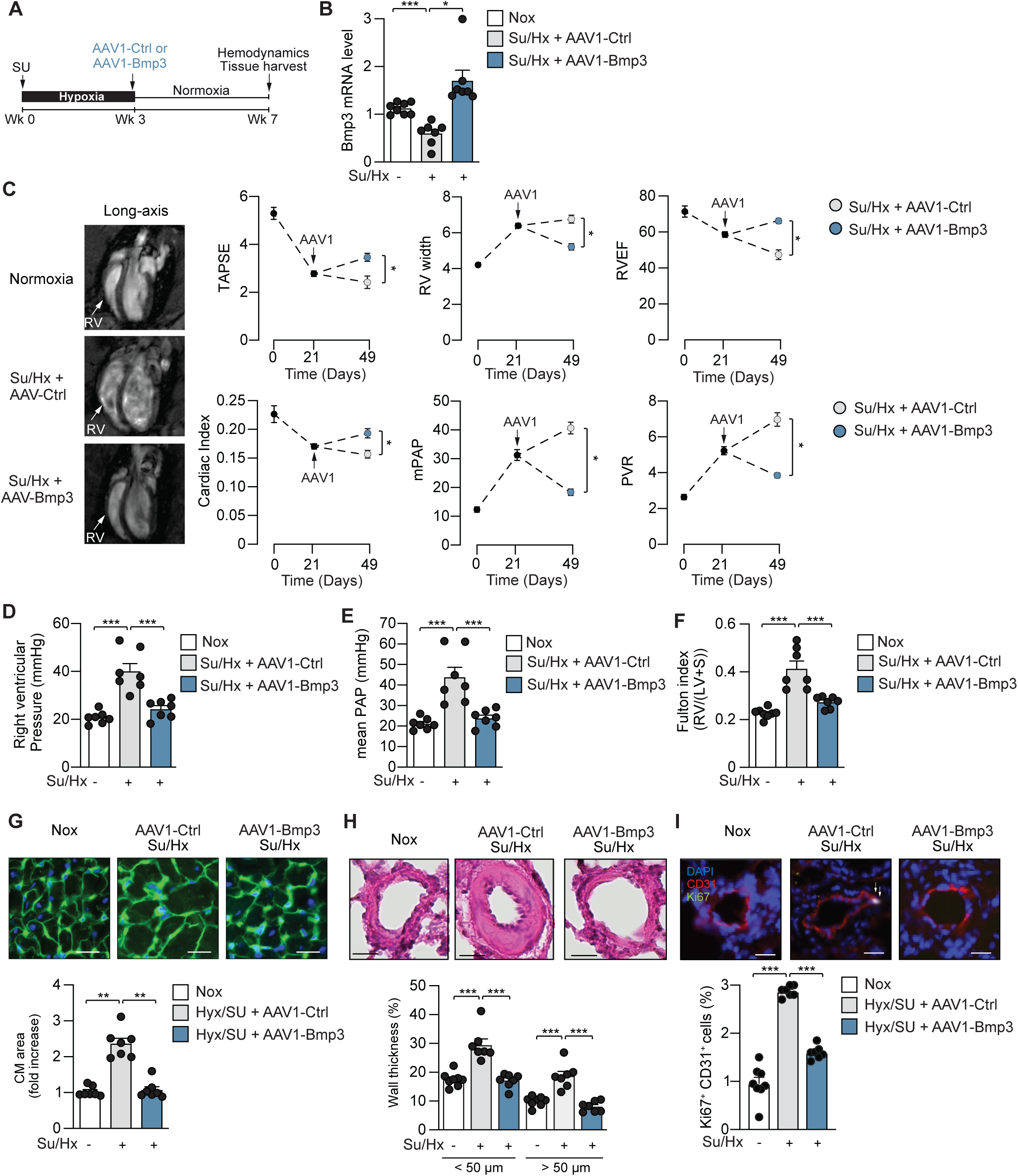
A lung-specific overexpression of BMP3 preserves the cardiac function and reverses pulmonary vascular remodeling in a severe model of PAH in rats. **A,** Design of the study. Rats received a single injection of Sugen (20 mg/kg) and were then exposed to three weeks of chronic hypoxia. Rats were then randomly assigned to receive AAV1-Ctrl or AAV1-Bmp3 at week 3 before being returned under normoxic conditions. The endpoint for hemodynamic measurements and sacrifice was at week 7. Cardiac MRI measurements were performed at weeks 0, 3 and 7. **B,** Bmp3 mRNA expression in lung homogenates from the indicated groups. n = 7-8 rats/group. **C**, Cardiac MRI parameters of the indicated conditions. n = 3-5 rats/group. **D**, RVSP of the indicated groups. n = 7 rats/group. **E**, mean pulmonary artery pressure (mPAP) of the indicated groups. n = 7 rats/group. **F**, Fulton index of the indicated groups. n = 7-8 rats/group. **G**, (Upper panel) Representative H&E-stained pulmonary artery sections from the indicated groups. Scale bar: 50 μm. (Lower panel) Percentage of arterial wall thickness in relation to cross-sectional diameter. n = 7-8 rats/group. **H**, (Upper panel) Representative WGA-staining of RV sections. Scale bar: 50 μm. (Lower panel) Quantitative analysis. n = 7 rats/group. **I**, Ki-67/CD31 positive cells in normoxic and Su/Hx lungs treated with AAV1-Ctrl or AAV1-Bmp3. n= 7-8 rats/group. Scale bar: 25 μm. ** P < 0.01; *** P < 0.001, by 1-way ANOVA or two-tailed *t* test. RV: Right ventricle, Su/Hx: Sugen/Hypoxia, CM: Cardiomyocyte. TAPSE: tricuspid annular plane systolic excursion, RV width: right ventricular width, RVEF: right ventricular ejection fraction, mPAP: mean Pulmonary Arterial Pressure, PVR: Pulmonary Vascular Resistance.

Consistent with our findings in the Su/Hx model in mice, BMP3 overexpression reduced RV and cardiomyocyte hypertrophy (Figure 6G), and morphometric analysis of distal pulmonary arteries demonstrated a significant decrease in the wall thickness of AAV1-Bmp3-treated rats (Figure 6H). Additionally, AAV1-mediated BMP3 overexpression attenuated endothelial cell proliferation as shown by reduced CD31+/Ki-67+ cells in the severe Su/Hx rat model of PAH (Figure 6I). These results support the therapeutic potential of BMP3 overexpression in treating severe pulmonary vascular disease.

### BMP3 Overexpression Restores the Balance Between the TGF-β/Smad2,3 and the BMP/Smad1,5,8 Signaling Pathways in PAH

Next, we sought to investigate the mechanisms by which BMP3 exerts therapeutic effects in PAH and its impact on the balance between the pro-proliferative TGF-β/SMAD2/3 pathway and the antiproliferative BMP/SMAD1.5/8 pathway. Using hPAEC cells transduced with a BMP response element (BRE)-GFP construct, we observed that BMP3 treatment significantly increased GFP expression, indicating activation of the BMP/Smad1,5,8 pathway (Figure 7A). In contrast, TGF-β-stimulation activated the Smad2/3 response elements (SRE)-GFP signal, an effect that was significantly inhibited by recombinant BMP3 (Figure 7B).

**Figure 7:**
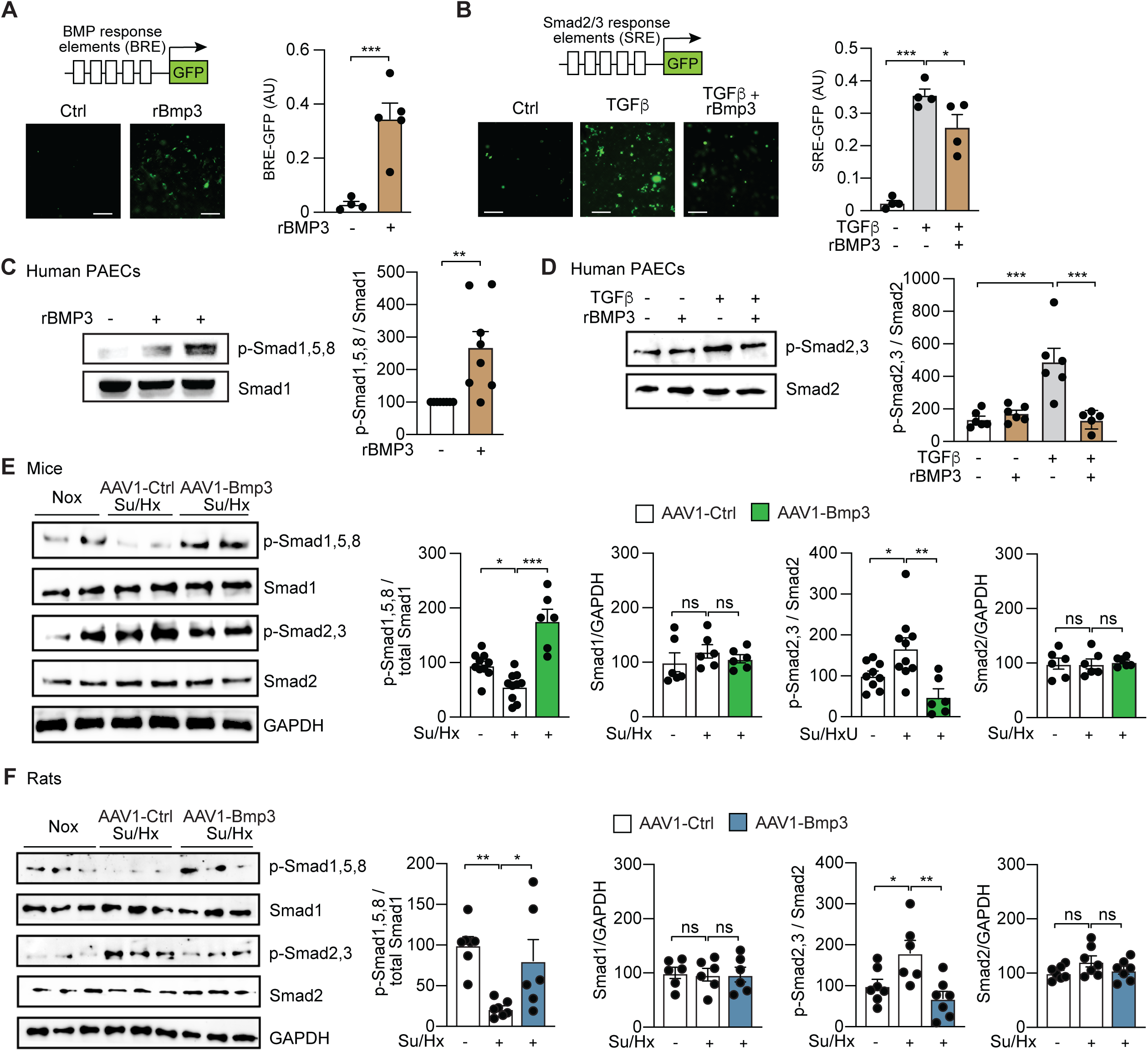
BMP3 regulates the TGF-β/Smad2,3 and the BMP/Smad1,5,8 pathways. **A**, GFP fluorescent intensity in hPAECs transduced with a lentivirus expressing BMP response elements (BRE)-GFP in the presence or absence of rBMP3 (100 ng/ml). n = 5 experiments performed in triplicate. **A**, GFP fluorescent intensity in hPAECs transduced with a lentivirus expressing Smad2/3 response elements (SRE)-GFP in the presence or absence of TGF-β1 (5 ng/ml) and rBMP3 (100 ng/ml). n = 5 experiments performed in triplicate. **C**, (Left) Phosphorylated Smad1/5/8 and total Smad1 protein expression in hPAECs in the presence or absence of rBMP3 (100 ng/ml). (Right) Quantification of the data. n = 8. **D**, (Left) Phosphorylated Smad2/3 and total Smad2 protein expression in hPAECs in the presence or absence of absence of TGF-β1 (5 ng/ml) and rBMP3 (100 ng/ml). (Right) Quantification of the data. n = 6. **E**, (Left) Phosphorylated Smad1/5/8, total Smad1, phosphorylated Smad2/3, total Smad2, and GAPDH protein expression in lung homogenates from healthy and Su/Hx mice treated with AAV1-Ctrl or AAV1-Bmp3. (Right) Quantification of the data. n = 6-10 mice/group. **F**, (Left) Phosphorylated Smad1/5/8, Smad1, phosphorylated Smad2/3, Smad2, and GAPDH protein expression in lung homogenates from healthy and Su/Hx rats treated with AAV1-Ctrl or AAV1-Bmp3. (Right) Quantification of the data. n = 6-7 rats/ group. * P < 0.05; ** P < 0.01; *** P < 0.001; ns: not significant by two-tailed *t* test or 1-way ANOVA.

Western blot analyses confirmed that rBMP3 treatment induced phosphorylation of Smad1/5/8 (Figure 7C), while simultaneously blocking TGF-β-induced phosphorylation of Smad2/3 in hPAECs (Figure 7D). In the lungs of AAV1-control-treated PH mice, we found a marked downregulation of phospho-Smad1/5/8, and a concurrent upregulation of phospho-Smad2/3 (Figure 7E). Consistent with our *in vitro* findings, BMP3 overexpression reversed this imbalance by restoring Smad1/5/8 phosphorylation and suppressing Smad2/3 phosphorylation (Figure 7E). These results were validated in Su/Hx-exposed rats, where AAV1-BMP3 treatment led to rebalancing of SMAD signaling with enhanced Smad1/5/8 phosphorylation and reduced Smad2/3 phosphorylation (Figure 7F).

Given that Smads translocate from the cytoplasm to the nucleus and regulate the transcription of target genes, we next performed RNA sequencing analysis to identify the gene targets of Smad signaling and downstream of BMP3 regulation. HPAECs were treated with conditioned medium from BMP3-silenced or BMP3-upregulated PASMCs, and total RNA was subsequently extracted for RNA-seq analyses (Figure 8A). KEGG pathway enrichment analysis revealed that cell cycle-related pathways were significantly enriched in hPAECs treated with CoM from Bmp3 depleted PASMCs (Figure 8B). Similarly, KEGG pathway analysis of hPAECs exposed to coM from BMP3 overexpressing PASMCs confirmed the involvement of cell cycle gene regulation (Figure 8C). Notably, a set of cell cycle regulators such as cyclins (Ccnb1, Ccne1, Ccna2, Ccnb2) and cyclin-dependent kinase (Cdk1), as well as mitotic checkpoint and cell division genes (Cdc19, Cdc46, Cdc20, and Plk1) were among the top differentially expressed genes. These genes expression levels were confirmed at the mRNA levels in PAECs treated with PASMCs’ CoM (Figure 8D). The in vivo Sugen/Hypoxia conditions induced a marked upregulation of these cell cycle-related genes in the lungs, an effect that was markedly inhibited by BMP3 overexpression (Figure 8E). Collectively, these results demonstrate that BMP3 overexpression modulates PAEC cell cycle activity by restoring the balance between the TGF-β/Smad2,3 and the BMP/Smad1,5,8 pathways, thereby suppressing aberrant activation of cell cycle-related genes in the context of PAH.

**Figure 8:**
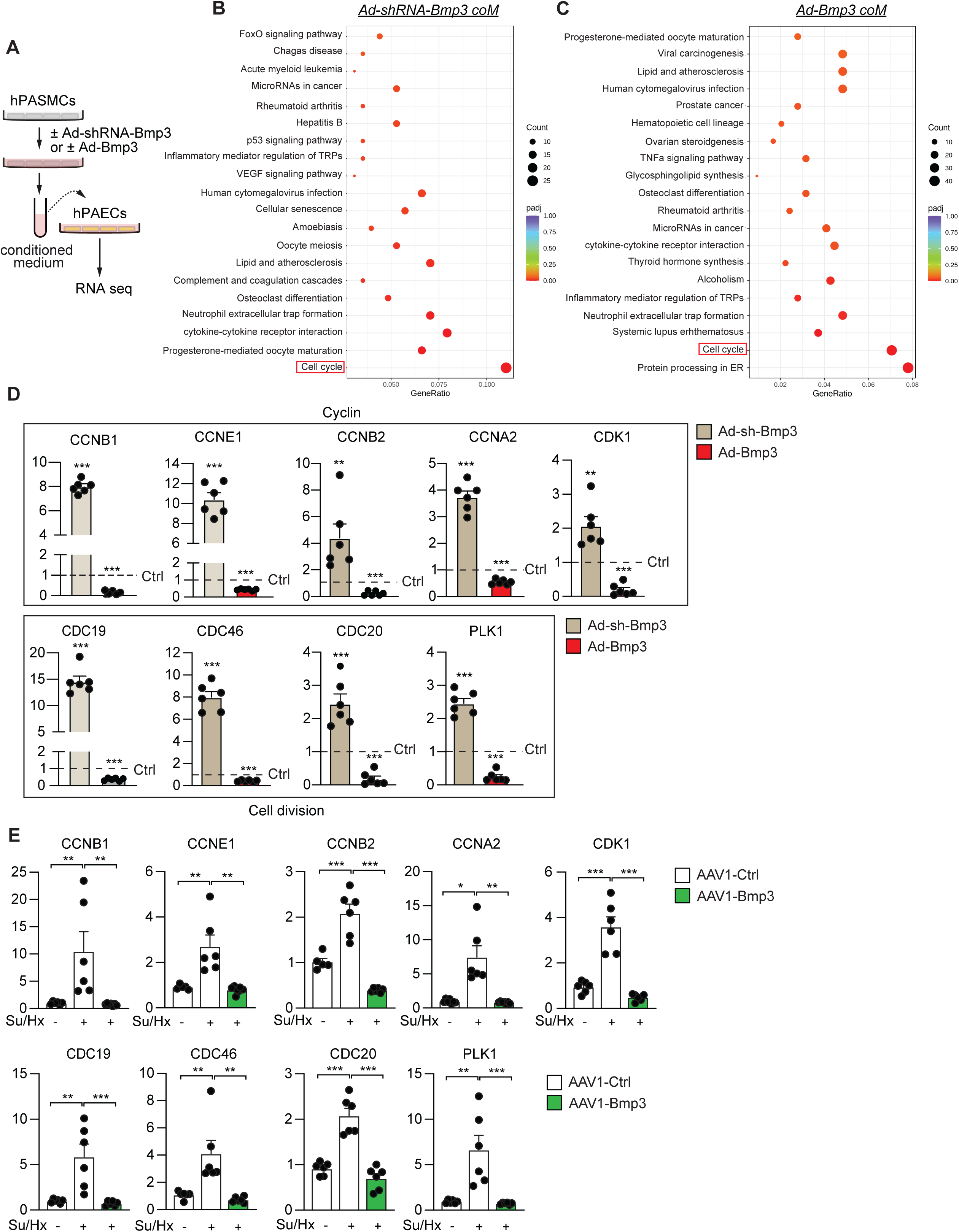
BMP3 regulates cell cycle gene expression in hPAECs. **A**, Experimental scheme to identify the pathways regulated by BMP3. PASMCs were infected with Ad-shRNA-Bmp3, Ad-Ctrl, or Ad-Bmp3. The conditioned medium (coM) from hPASMCs was then added to hPAECs. 48 hours later, RNA was extracted and RNA sequencing was performed. **B**, Scatter plot of top KEGG pathways enriched by differentially expressed genes in hPAECs treated with Ad-control vs Ad-shRNA-Bmp3 coM (n = 4 per group). **C**, Scatter plot of top KEGG pathways enriched by differentially expressed genes in hPAECs treated with Ad-control vs Ad-Bmp3 coM (n = 4 per group). **E**, PCR analysis of ccnb1, ccne1, ccnb2, ccna2, cdk1, cdc19, cdc46, cdc20, and Plk1 mRNA levels in hPAECs treated with Ad-control, Ad-shRNA-Bmp3, or Ad-Bmp3 coM (n = 6 per group). **C**, Pulmonary ccnb1, ccne1, ccnb2, ccna2, cdk1, cdc19, cdc46, cdc20, and Plk1 mRNA levels in lungs from the indicated groups (n = 5-7 mice per group). * P < 0.05; ** P < 0.01; *** P < 0.001; ns: not significant by two-tailed *t* test or 1-way ANOVA.

## DISCUSSION

In this study, we identified BMP3 as a novel regulator of pulmonary vascular homeostasis. BMP3 is predominantly produced by PASMCs and is downregulated in PAH. We demonstrated that PASMC-derived BMP3 acts as a paracrine factor inhibiting the proliferation and migration of PAEC. The loss of BMP3, either globally or specifically in SMCs in mice, worsens PAH in middle-aged mice, highlighting its protective role. Therapeutic BMP3 overexpression, using a recombinant protein or an AAV-mediated lung gene delivery, prevented and reversed PAH in rodent models. Mechanistically, this effect was associated with the restoration of the balance between the proliferative TGF-β/Smad2,3 and the antiproliferative BMP/Smad1,5,8 pathways. Together, our findings reveal an unprecedented paracrine role of BMP3 in pulmonary vasculature and establish that BMP3 is a new promising therapeutic target for the treatment of PAH.

Global or SMC–specific Bmp3-deficient young adult mice did not exhibit any obvious phenotype under baseline conditions or following exposure to the Su/Hx, suggesting age-dependent effects of BMP3 deficiency in PAH progression. Interestingly, Bmp3-deficient mice showed no significant changes in RV hypertrophy and pulmonary vascular morphology when compared to WT controls, indicating that BMP3 deficiency alone is insufficient to drive pathology in mice. In contrast, middle-aged (9-12-month-old) global or SMC–specific Bmp3-deficient mice exhibited exacerbated pulmonary vascular remodeling and RV dysfunction in response to Su/Hx conditions. These observed findings suggest that BMP3 plays a more critical protective role during aging and under stress conditions such as VEGF inhibition and chronic hypoxia. The discrepancy between the young and middle-aged Bmp3-deficient mice might be attributed to the presence of a transient compensatory mechanism in younger mice that likely diminishes with age. This compensatory mechanism may progressively disappear once a critical stage of pulmonary vascular remodeling has been reached.

Previous studies have shown that BMP3 co-localizes with Activin A Receptor Type IIB (ActRIIB) at the plasma membrane in a colon cancer cell line (KM12), and that the knockdown of endogenous ActRIIB was associated with a reduction in the growth-suppressive effect of BMP3 in bone marrow stromal cells.^22,28^ During Xenopus embryogenesis, BMP3 has been reported to antagonize and inhibit both Activin and BMP4 signaling, in part by interfering with activin binding to ActRIIB, thus leading to decreased Smad2 activation and blocking activin signaling transduction.^29^ Importantly, BMP3 binds to ActRIIB in a non-competitive state, as excess activin cannot displace BMP3, resulting in a receptor complex that fails to activate Smad2 and transduce the signal.^29^ It remains unclear whether BMP3 signals in PAECs by binding to Activin A Receptor or by binding to BMPR or a combination of both.

Treatment of hPAECs with rBMP3 resulted in increased Smad1/5/8 phosphorylation, along with reduction of TGF-β-induced Smad2/3 phosphorylation, indicating a dominant shift to BMP downstream signaling. In line with these in vitro findings, BMP3 overexpression restored Smad1/5/8 phosphorylation and decreased Smad2/3 phosphorylation in the lungs of PAH-diseased mice and rats, suggesting that BMP3 restores the balance between the TGF-β/Smad2,3 and the BMP/Smad1,5,8 pathways. While our results strongly indicate significant effects of BMP3 in modulating BMP/TGFβ pathways, it is crucial to acknowledge that the observed improvement in RV function and pulmonary vascular remodeling may not be entirely attributable to this mechanism. It remains possible that the observed therapeutic effects may be attributed to BMP3’s effects on additional downstream pathways. Thus, identifying and characterizing these potential pathways will require further investigation.

The human genome encodes more than 20 BMP ligands that signal through homodimeric or heterodimeric forms.^30^ However, only a limited number of BMPs have been investigated in the context of PAH. BMP4, a ligand that is secreted by endothelial cells in response to hypoxia, promotes the proliferation and migration of vascular SMCs.^14,31^ Consistently, BMP4 KO mice are protected from developing hypoxia-induced PH and exhibit reduced pulmonary vascular cell proliferation and vascular remodeling.^14,31^ Interestingly, BMP2, a homologous protein of BMP4, exerts opposing roles in PAH.^32^ Genetic deletion of BMP2 in mice exacerbates hypoxia-induced PH^32^, while upon administration of exogenous recombinant BMP2 suppresses PASMC proliferation and migration, suggesting a protective role of BMP2 in PH disease pregression.^16^ BMP9 has been shown to exert dual and context-dependent effects in PAH.^18,33^ Similarly, BMP-11 (also known as GDF11) is significantly upregulated in PAH, and its specific deletion in endothelial cells attenuates hypoxia-induced PAH and pulmonary vascular remodeling in mice.^34^ Despite intensive efforts to understand the role of BMP receptors in PAH, only a few BMP ligands have been studied in the context of PAH.^17,35,36^ Further investigating the roles of other BMP ligands will contribute to a better understanding of the pathogenesis of PAH and may uncover and advance novel target therapies for this fatal disease.

Sotatercept, a fusion protein consisting of the extracellular domain of the activin receptor type IIA fused to the Fc portion of human IgG1, is a novel therapeutic agent used in the treatment of PAH.^37^ Sotatercept acts as a ligand trap for activins and growth differentiation factors (GDFs), members of the TGF-β superfamily ligands. By trapping excess activins, Sotatercept shifts the signaling balance toward the BMP-dominant pathway, promoting anti-proliferative and anti-inflammatory processes in pulmonary vascular cells. Blocking GDF8/11- and activin-mediated SMAD2/3 activation in vascular cells and activating the SMAD1/5/8 pathway, by combining Sotatercept and BMP3 overexpression, may offer a synergistic approach for attenuating the development of PAH.

A limitation of this study is the lack of direct measurement of BMP3 protein expression in the lungs of PAH-diseased animals and in pulmonary vascular cells derived from patients with PAH. Despite testing of different commercially available BMP3 antibodies, in cells upon the modulation of BMP3 and tissues from BMP3-deficient mice, none of the antibodies showed a specific band for BMP3 in western blotting (data not shown). The development of specific antibodies for BMP3 is essential to accurately detect and quantify BMP3 protein levels both in circulating blood and pulmonary tissues. This would significantly enhance our ability to investigate BMP3 dynamics in PAH pathogenesis and could support the use of BMP3 as a biomarker or therapeutic target in future studies in PAH.

It is important to note that the SMC-specific deletion of BMP3 mirrored the effects of global BMP3 deletion, indicating that blocking the production of BMP3 in SMCs is sufficient to halt its protective paracrine signaling in PAH. BMP ligands have been shown to mediate paracrine signaling across different tissues and physiological contexts.^38^ The paracrine effects of BMP ligands are integral to various biological processes, including tissue development, regeneration, and disease progression. In the intestine, BMPs are secreted by niche cells to maintain stem cell quiescence or promote differentiation in nearby progenitors.^39^ BMP2/4 act in a paracrine manner to hematopoietic progenitors, leading to megakaryocytic lineage phenotype of the leukemia.^40^ BMP4 negatively regulates C19 steroid synthesis through its paracrine signaling in the human adrenal glands.^41^ BMP3, secreted from PASMC, may circulate in the blood and act on distant organs or vascular beds.

BMP3 is known to play important roles in different organs and biological processes. Mice with increased BMP3 levels in bone exhibit delayed endochondral ossification.^21,42^ Expression of exogenous BMP3 suppresses the proliferation and migration of colon cancer cell lines, and inhibits tumor growth in immunodeficient mice.^43^ BMP3 up-regulation protects against bleomycin-induced pulmonary fibrosis in mice.^44^ Additionally, BMP3 inhibits TGFβ-induced myofibroblast differentiation during embryonic cornea wound healing.^45^ Recombinant BMP3 has also emerged as a promising novel therapeutic strategy for kidney fibrosis.^46^ Importantly, to date, no deleterious effects following BMP3 overexpression have been reported. Given its wide roles, potential off-target effects in other organs may result from BMP3 overexpression; however, in the context of a targeted lung therapy using intratracheal inhalation, the side effects are reduced.

Taken together, our study identifies BMP3 as a critical paracrine mediator between pulmonary vascular cells by suppressing hPAEC dysfunction, thereby establishing BMP3 as a promising therapeutic target in PAH.

## Acknowledgments

The authors thank Dr. Olympia Bikou, Clemens Eisenacher, and Gaia Chen for technical support, and gratefully acknowledge the Histology resources and services provided by the FBRI Histology Core Facility at Virginia Tech Carilion, and the BioMedical Engineering and Imaging Institute (BMEII) at the Icahn School of Medicine at Mount Sinai (ISMMS).

## SUPPLEMENTAL MATERIAL

## EXPANDED METHODS

### Proliferation Assay

To assess cell proliferation, PAECs or PASMCs were seeded in 96-well plates at 70% confluence before being treated with recombinant BMP3 or conditioned media. 48 hours later, a BrdU labeling solution was added to each well and incubated for 4-16h, following the manufacturer’s instructions. Subsequently, cells were fixed using the provided fixative. The cells were then incubated with anti-BrdU antibody. After thorough washing, BrdU incorporation was detected using an appropriate substrate for colorimetric measurement, and the signal was quantified using a plate reader (GloMax). To assess the direct effect of BMP3 on PAEC and PASMC proliferation, cells were treated with human recombinant BMP3 (rBMP3, 100 ng/mL; PeproTech) following the same experimental protocol.

### Wound Healing Assay

PAECs or PASMCs were seeded in a 12-well plate at a high density (80%-90% confluence) and incubated with VascuLife VEGF Endothelial Medium (LifeLine, LS-1029) containing 5% FBS. After reaching a confluent monolayer, scratching was performed and examined under a microscope. After removing any detached cells by washing, 500 uL of 0.1% FBS fresh culture medium and 500 uL of CoM collected from PASMC were added to the wells. Afterwards, plates were incubated and wells were then photographed at different time points (0 h, 24 h, and 48 h). To evaluate the direct effect of BMP3 on PAEC and PASMC migration, cells were treated with rBMP3 following the same experimental protocol. ImageJ software was used to measure the width of the scratch at different time points. To calculate the percentage of wound area we used the formula: Wound area (%) = (Initial Wound Width−Final Wound Width/Initial Wound Width) ×100.

### Bmp3 deficient mice

The mice were housed under a 12-h light/12-h dark cycle with ad libitum access to food and water. All procedures were conducted in accordance with the guidelines set by the Institutional Animal Care and Use Committee (IACUC) of Virginia Tech.

Conditional BMP3 KO mice (Bmp3^flox/flox^) were originally generated in the lab of Dr. Vicki Rosen (Harvard School of Dental Medicine). To generate global BMP3 KO mice, we crossed Bmp3^flox/flox^ mice with UBC-CreER^T2^. Littermates homozygous for Bmp3^flox/flox^ but lacking the Cre allele served as controls. Mice (8-10 weeks old) were administered daily intraperitoneal (i.p.) injections of tamoxifen (TAM, 40mg/Kg; Sigma) for five consecutive days.^23^ To generate SMC-specific BMP3 KO mice, we crossed Bmp3^flox/flox^ mice with SM22-Cre mice. Littermates homozygous for Bmp3^flox/flox^ but lacking the Cre allele served as controls. Young (8 weeks old) and middle-aged (9-12 months-old) mice were exposed to chronic hypoxia (10% O₂) for four weeks in a hypoxia chamber, with SU5416 (Sugen, 20mg/kg; MedChemExpress) being subcutaneously injected weekly into the mice (during 3 weeks). Parallel groups of knockout and control mice were maintained under normoxic conditions (20% O₂) as controls. Following the four-week exposure period, hemodynamic measurements were performed using pressure-volume (PV) loop measurements.

### Recombinant BMP3 Treatment in Mice

In the prevention study, every 3 days, mice intraperitoneally received a dose of recombinant BMP3 (0.3 mg/kg, Peprotech) or saline (as control) and were maintained in hypoxia for 3 weeks with a weekly injection of Sugen. Hemodynamic measurements and tissues harvest were performed at week 3.

In the therapeutic study, mice were first exposed to hypoxia for two weeks with a weekly injection of Sugen, before receiving rBMP3 (0.3 mg/kg, Peprotech) or saline injection every 3 days for an additional two weeks. Parallel groups of mice were maintained under normoxic conditions (20% O₂) as controls. At the end of the exposure period, hemodynamic analysis was performed using PV-loop measurements.

### AAV1-Mediated BMP3 Overexpression

In the prevention study: Wild-type C57BL/6 mice (8-week-old) were randomized to intratracheally receive either aerosolized AAV1-Bmp3 or AAV1 encoding luciferase as control (2×10^11^ genome copies per mouse) two weeks before being exposed to hypoxia for 4 weeks with a weekly injection of Sugen. Hemodynamics and morphometric measurements were performed 6 weeks after AAVs delivery. In the therapeutic study, mice were exposed to Sugen/hypoxia for 2 weeks and were then randomly assigned to receive AAV1-Bmp3 or AAV1-Ctrl treatment. Hemodynamics and morphometric measurements were performed 4 weeks after AAVs delivery. Parallel groups of mice were maintained under normoxic conditions (20% O₂) as controls. AAV1-CMV-Luciferase (control) or AAV1-CMV-mBMP3 were provided by the Penn Vector Core Gene Therapy Program (University of Pennsylvania).

### Pig PH model

Lung tissues were obtained from PH-diseased Yorkshire pigs. Pulmonary hypertension was induced using pulmonary vein banding (postcapillary Pulmonary Hypertension) as described previously.^24,25^ Briefly, 10-12 kg pigs were sedated with Telazol, intubated and anesthetized with isoflurane. Thereafter, basic cardiac functional measurements were performed (echocardiography and right heart catheterization). PH was surgically induced through banding of the superior and inferior pulmonary veins. The pulmonary veins were banded and the success of the procedure was confirmed by echocardiography, with the application of pulsed wave Doppler. Four months after pulmonary vein banding, invasive and non-invasive cardiac function measurements were performed and the animals were sacrificed. Lungs were perfused with 0.9% saline solution before further processing. Lung tissues were snap-frozen for further molecular analysis. All samples used in this study are from the lower lung lobes. The hemodynamic data of the animals are shown in Table S3.

### Magnetic Resonance Imaging (MRI) Analysis

Cardiac magnetic resonance imaging (MRI) was performed at three time points: baseline, 3 weeks, and 7 weeks. For MRI acquisition, rats were positioned on the instrumentation panel bed and placed into a Bruker 7T small animal MRI system (Bruker AXS, Inc., Madison, WI). Respiration-gated MRI scans were obtained in both short-axis and long-axis views. Images were acquired with a slice thickness of 1.5 mm to ensure complete coverage of the biventricular anatomy. All DICOM images were imported into SEGMENT MRI software and analyzed by a trained analyst. The short-axis and long-axis CINE series were evaluated for cardiac dimensions, volumes, ejection fraction, myocardial mass, regional function, and chamber indices using semi-automated contour segmentation. Scans were anonymized and analyzed in batches by an analyst who was blinded to the hemodynamic data at the time of evaluation. Stroke volume (SV) was calculated as the difference between end-diastolic volume (EDV) and end-systolic volume (ESV). Ejection fraction (EF) was computed using the formula: EF = (SV / EDV) × 100. Cardiac output (CO) was calculated from MRI data using the formula: CO = SV × HR, where HR is the average heart rate recorded during the scan. Cardiac index (CI) was calculated by dividing the CO by the body surface area (BSA). The total rat’s BSA of each animal was calculated using Meeh’s formula. Tricuspid annular plane systolic excursion (TAPSE) was assessed using the four-chamber MRI views. The maximum distances between the lateral tricuspid annulus and the right ventricular apex were measured at end-systole and end-diastole. TAPSE was defined as the difference between these two measurements. Myocardial mass was quantified by acquiring a volumetric dataset of the heart. The epicardial and endocardial contours of the ventricle were manually delineated, and the segmented myocardial volume was multiplied by the specific gravity of myocardial tissue (1.05 g/cm³) to calculate ventricular mass. All ventricular volume and mass measurements were indexed to body surface area.

### Hemodynamics Analysis

Mice and rats were anesthetized with isoflurane and placed on a heated platform for temperature regulation. A 1.4F or 1.9F pressure-volume catheter was inserted into the right ventricle or pulmonary artery. Right ventricular systolic pressure (RVSP) and mean pulmonary arterial pressure (mPAP) were measured using a pressure-volume (PV) loop system (ADInstruments) to continuously record the pressure waveform during the cardiac cycle. The animals were monitored throughout the procedure for anesthesia depth and recovery.

### Tissues Harvest

After completing the hemodynamic measurements, mice were perfused with phosphate-buffered saline (PBS) to remove blood. The heart and lungs were then harvested for further analysis. The Fulton Index, which is the ratio of the right ventricle (RV) weight to the combined weight of the left ventricle and septum (LV+S), was calculated as an indicator of right ventricular hypertrophy. Pieces of the right ventricle (RV), left ventricle (LV), and lungs were collected for RNA and protein extraction. Additionally, a portion of the RV or lung tissue was placed in optimal cutting temperature (OCT) compound for histopathological analysis.

### RNA Extraction

Total RNA was extracted, from cells or grounded tissues, using the TRIzol reagent (Invitrogen), following the manufacturer’s protocol. Briefly, samples were homogenized in TRIzol, and after phase separation, the aqueous phase containing RNA was carefully collected. RNA was then precipitated using isopropanol, washed with 75% ethanol, and resuspended in RNase-free water. To remove any residual DNA, RNA samples were treated with DNase I (Thermo Scientific) following the manufacturer’s instructions. RNA quantity and quality were assessed using a NanoDrop spectrophotometer (Thermo Scientific), and samples were stored at -80°C until further analysis.

### Reverse Transcription

For reverse transcription, 500 ng of total RNA was used as the template for cDNA synthesis using the High-Capacity cDNA Reverse Transcription Kit (Applied Biosystems), following the manufacturer’s protocol. The reaction was performed at 25°C for 10 min, 37°C for 120 min, and 85°C for 5 min to inactivate the reverse transcriptase. The resulting cDNA was stored at -20°C for downstream quantitative PCR (qPCR) analysis.

### Real-time quantitative PCR

Quantitative PCR (qPCR) was performed using SYBR Green Master Mix (Quanta bio) and gene-specific primers for target genes. The reaction conditions included an initial denaturation at 95°C for 10 min, followed by 40 cycles of denaturation at 95°C for 15 sec, annealing at 60°C for 30 sec, and extension at 72°C for 30 sec. Melting curve analysis was performed after amplification to confirm primer specificity. Gene expression levels were normalized to housekeeping genes GAPDH, and the relative expression of target genes was calculated using the ΔΔCt method, where the Ct value of each gene in the experimental group was compared to the control group to determine fold changes in expression. Primers’ sequences are shown in Table S4 and S5.

### Protein Extraction

For protein extraction, cells or lung tissues were homogenized in RIPA buffer (Sigma-Aldrich) containing a phosphatase and protease inhibitor cocktail (Sigma-Aldrich). The samples were centrifuged at 12,000 × g for 20 min at 4°C to remove cellular debris, and the supernatants were collected for analysis. Protein concentration was quantified using the BCA protein assay kit (Pierce, ThermoFisher), following the manufacturer’s instructions. The resulting protein samples were stored at -80°C for future analysis.

### Western Blot Analysis

Protein samples (20-30 µg) were resolved by SDS-PAGE and transferred to a nitrocellulose membrane (Bio-Rad). The membrane was blocked with 5% Bovine Serum Albumin (BSA; Sigma-Aldrich) in Tris-buffered saline with 0.1% Tween 20 (TBST; Boston BioProducts) for 1 h at room temperature (RT). The membrane was incubated overnight at 4°C with primary antibodies specific for pSMAD1/5/9, pSMAD2/3, SMAD1, SMAD2, phospho-eNOS and eNOS (both from BD Biosciences), and GAPDH (Proteintech). After washing, the membrane was incubated with appropriate horseradish peroxidase (HRP)-conjugated secondary antibodies (Cell Signaling Technology) for 1h at RT. Protein bands were detected using the Bio-Rad ChemiDoc Touch Imaging System with SuperSignal™ West Pico PLUS Chemiluminescent Substrate (Thermo Scientific). Densitometric analysis of the bands was performed using Image Lab software (Bio-Rad), and relative protein expression was normalized to control protein levels for each sample.

### Hematoxylin Eosin (H&E) Staining

Cryosections (8 µm) were cut using a cryostat and mounted onto glass slides. The slides were then processed for H&E staining to assess the wall thickness of pulmonary arteries. Briefly, sections were air-dried and fixed in 4% paraformaldehyde for 10-15 min. After fixation, the sections were stained with Hematoxylin for 10 min, followed by differentiation in 1% acid alcohol and counterstaining with Eosin for 2 min. The slides were then dehydrated through a series of alcohols, cleared in xylene, and mounted with a coverslip. Images were captured using a light microscope, and arterial wall thickness was measured using ImageJ software. The ratio of the medial area to the total arterial area was calculated to quantify wall thickness and assess the degree of remodeling in the pulmonary arteries.

### Wheat Germ Agglutinin (WGA) staining

RV tissue sections (8 µm) were obtained from OCT-embedded frozen samples and mounted onto glass slides. The sections were fixed in 4% paraformaldehyde for 15 min at RT to preserve tissue morphology. After fixation, the sections were washed in PBS to remove excess fixative. WGA conjugated to Alexa Fluor (Thermo Fisher Scientific) was applied to the tissue sections at a concentration of 1 mg/mL and incubated in a humidified chamber at 37 for 90 min. WGA binds to sialic acid and N-acetylglucosamine residues on cell membranes, specifically highlighting the cell membrane architecture of cardiomyocytes. After incubation, the sections were washed in PBS to remove unbound WGA. To counterstain the nuclei, sections were incubated with DAPI (10 µg/mL; Invitrogen) for 5 min at RT. Finally, the sections were mounted with a coverslip using an anti-fade mounting medium (Vector Laboratories). Images were captured using a fluorescence microscope (Keyence), and the RV tissue morphology was analyzed for hypertrophy.

### Ki-67 staining

OCT-embedded frozen lung sections (8 µm) were fixed in 4% Formaldehyde and permeabilized with 0.15 % Triton X-100 for 15 min at RT. After blocking with normal donkey serum, sections were incubated overnight at 4°C with primary antibodies against Ki-67 (ab16667, Abcam) and CD31/PECAM-1 (AF3628, R&D Systems). Appropriate secondary antibodies, Alexa Fluor™ 555-conjugated goat anti-mouse IgG and Alexa Fluor® 647-conjugated donkey anti-goat IgG, were applied for detection. Nuclei were counterstained with DAPI, and slides were mounted with antifade medium before imaging by fluorescence microscopy (Keyence). Quantification was performed by counting Ki-67-positive nuclei within CD31-positive vascular regions.

### BRE-GFP and SRE-GFP

hPAECs were seeded in 96-well plates and cultured overnight under standard conditions. The following day, cells were transduced with either Smad2/3-TAG-Puro (LTV-0013-1S, LipExoGen) or BRE-TAG-Puro (LTV-0075-1S, LipExoGen) lentiviral reporters for 24 hours. After transduction, cells were serum-deprived for 4 hours, followed by stimulation with rBMP3 (100 ng/ml) TGF-β (5 ng/ml), or their combination for 12-24h. The GFP signal was quantified by fluorescence microscopy.

## SUPPLEMENTAL FIGURES AND FIGURE LEGENDS

**Figure S1:**
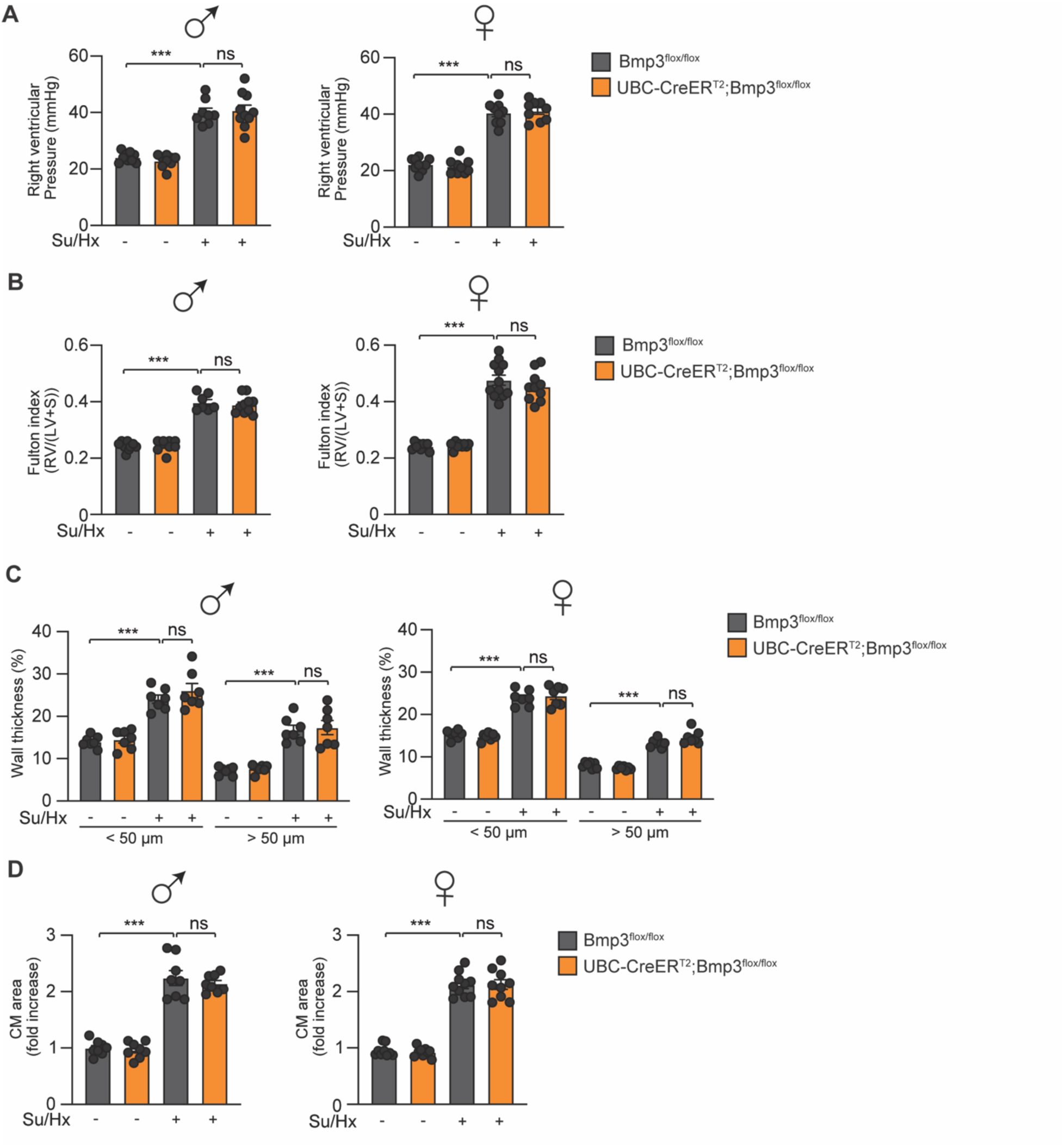
Global BMP3 deletion does not affect PAH in male and female young mice. **A,** Right ventricular systolic pressure (RVSP) of the indicated male (n = 8-10) and female (n = 9) groups. **B,** Fulton index of the indicated male (n = 9-10) and female (n = 9-12) groups. **C,** Percentage of arteries wall thickness of the indicated male (n = 7) and female (n = 7) groups. **D,** Quantitative analysis of cardiac myocytes size of the indicated male (n = 8) and female (n = 9-10) groups. *** P < 0.001; ns: not significant by 2-way ANOVA.

**Figure S2:**
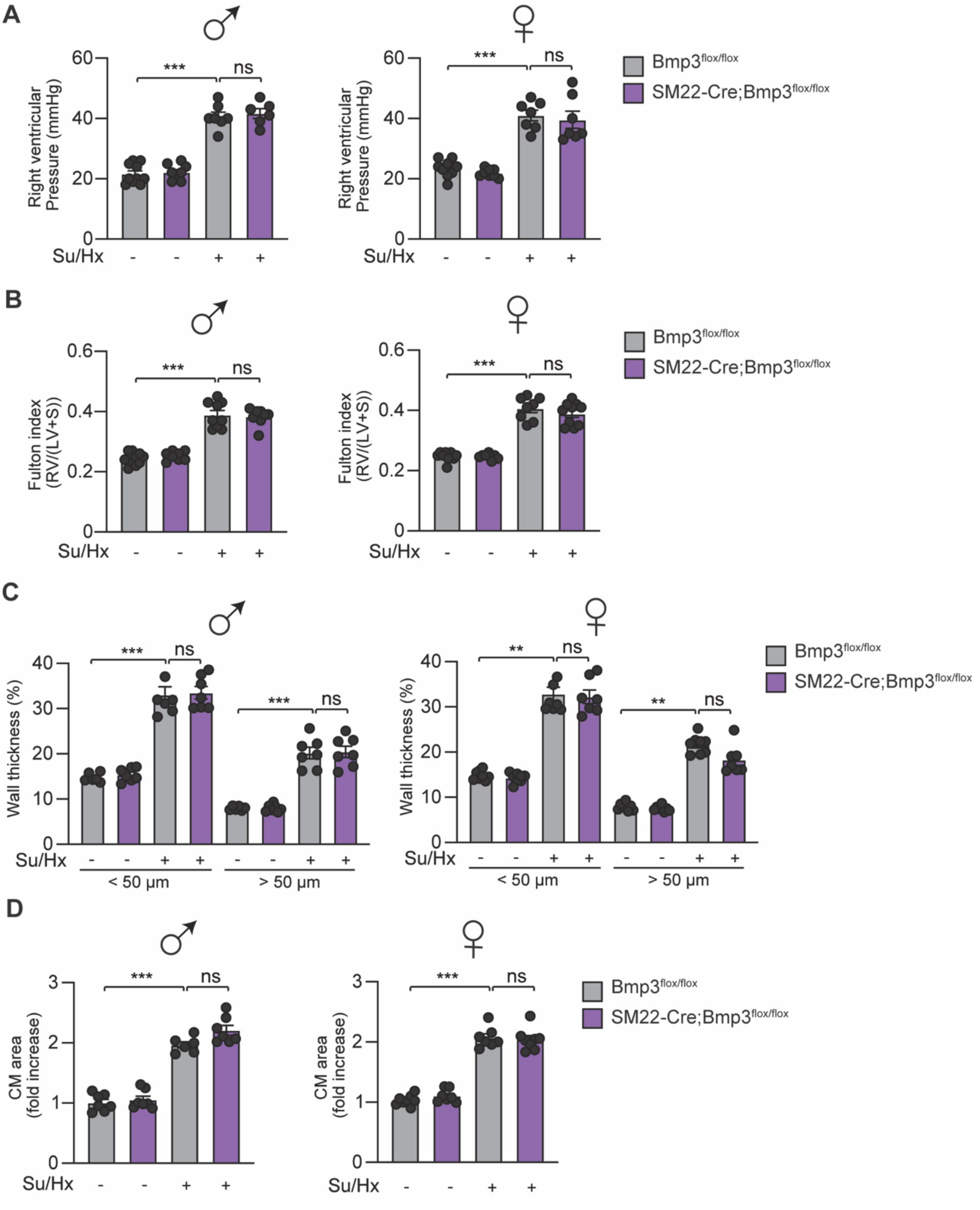
SMC-specific BMP3 deletion does not affect PAH in male and female young mice. **A,** Right ventricular systolic pressure (RVSP) of the indicated male (n = 6-10) and female (n = 7-9) groups. **B,** Fulton index of the indicated male (n = 8-10) and female (n = 8-10) groups. **C,** Percentage of arteries wall thickness of the indicated male (n = 6-7) and female (n = 7-8) groups **D,** Quantitative analysis of cardiac myocytes size of the indicated male (n = 6-7) and female (n = 6-7) groups. *** P < 0.001; ns: not significant by 2-way ANOVA.

**Figure S3:**
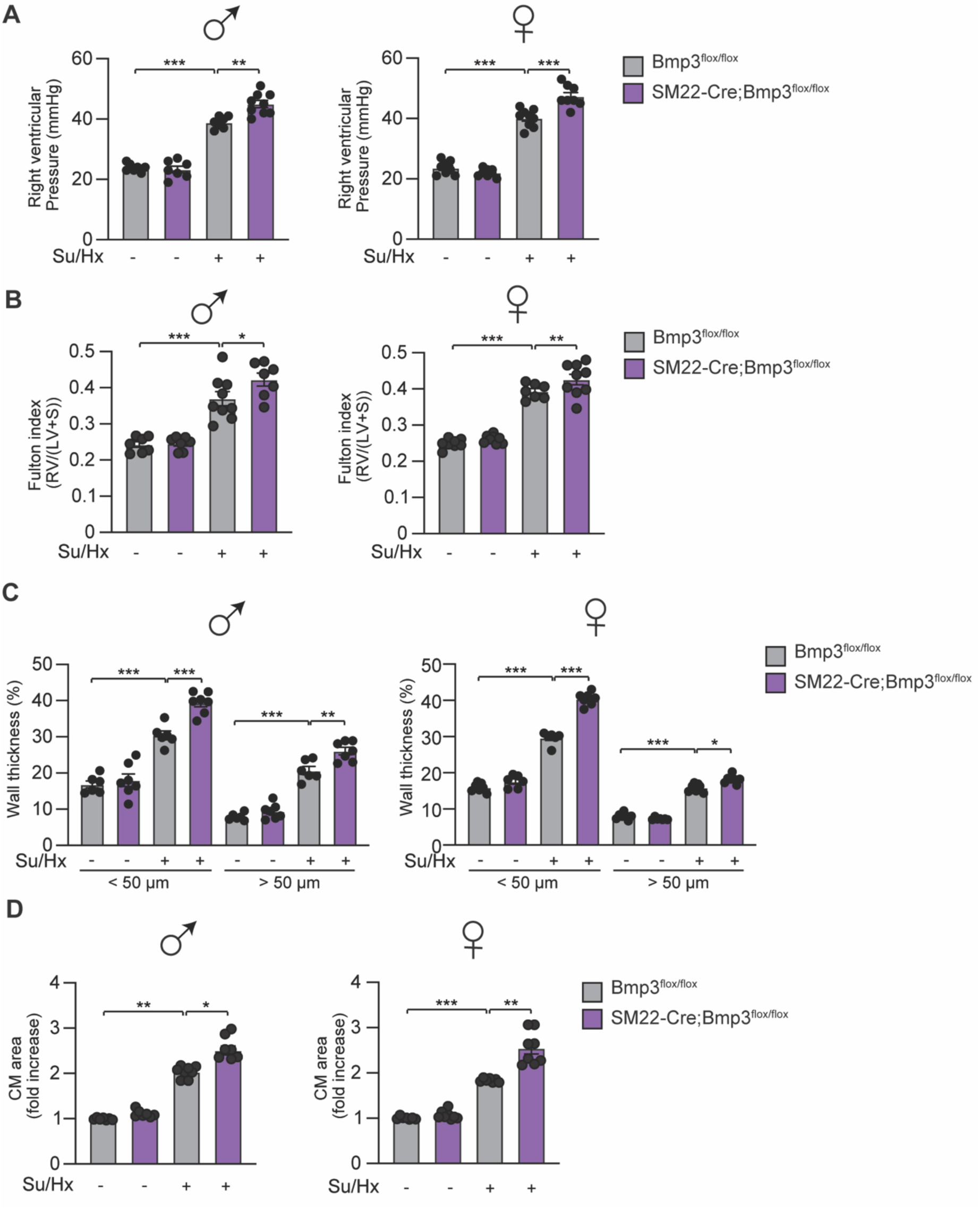
SMC-specific BMP3 deletion exacerbates PH in male and female mid­ dle-aged mice. **A,** Right ventricular systolic pressure (RVSP) of the indicated male (n = 7-9) and female (n = 8-9) groups. **B,** Fulton index of the indicated male (n = 7-9) and female (n = 8-9) groups. **C,** Percentage of arteries wall thickness of the indicated male (n = 6-7) and female (n = 6-7) groups. **D,** Quantitative analysis of cardiac myocytes size of the indicated male (n = 7-8) and female (n = 7-8) groups. • P < 0.05; •• P < 0.01; *** P < 0.001; ns: not significant by 2-way ANOVA.

**Figure S4:**
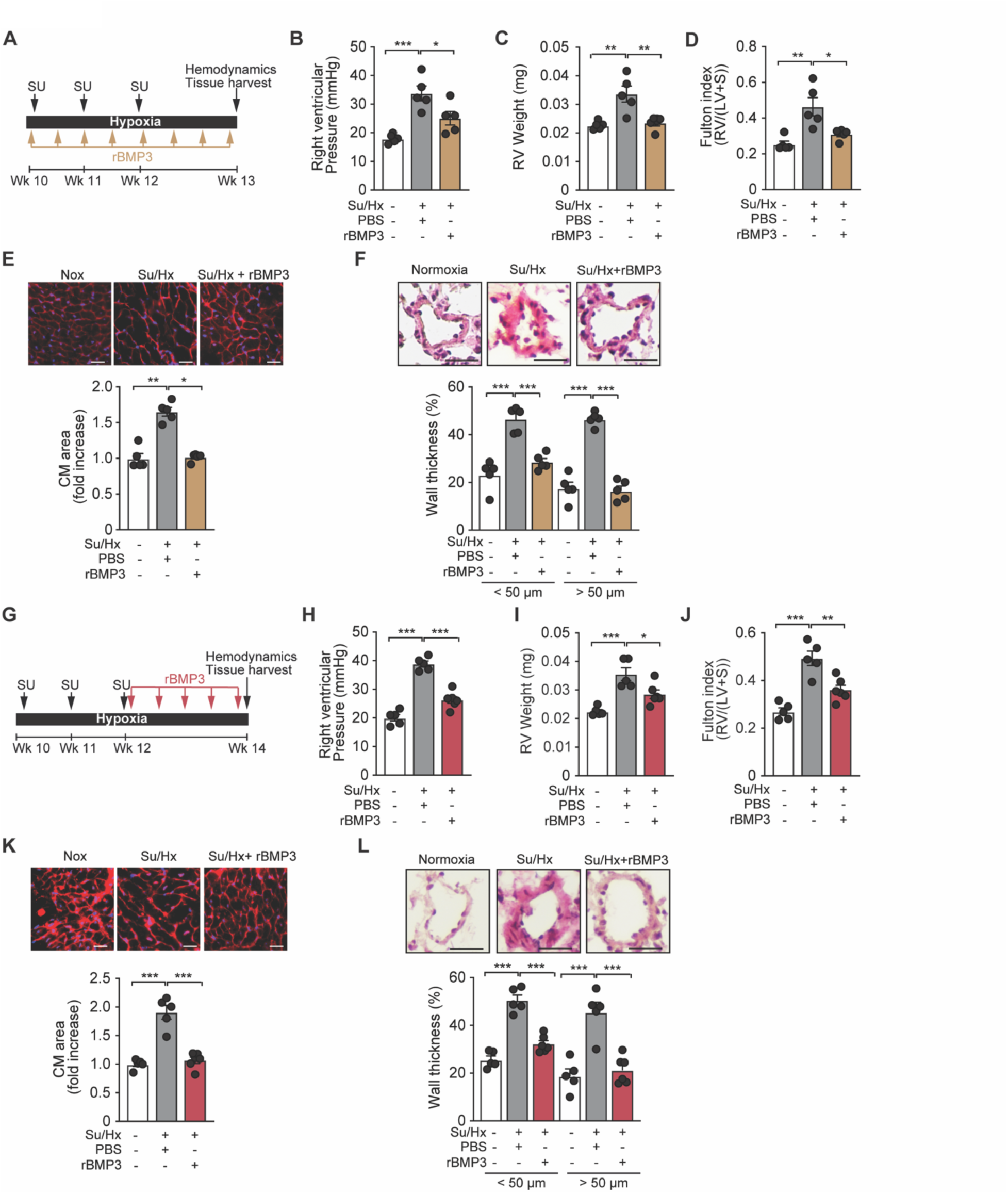
Recombinant BMP3 prevents and reverses PH and vascular remodeling in Sugen/Hypoxia-exposed mice. **A,** Design of the study. Ten-week-old C57B6 mice subcutaneously received 20 mg/kg of SU5416 (Su) and were then exposed to three weeks of chronic hypoxia. SU5416 was injected once a week during the next two weeks. The mice intraperitoneally received, every 3 days, a dose of recombinant BMP3 (rBMP3; 0.3 mg/kg) or saline as a control. The end point for hemodynamic measurements and sacrifice was at week 13. **B,** RVSP of the indicated groups. n = 5 mice/group. **C,** Right ventricular weight of the indicated groups. n = 5 mice/group. **D,** Fulton index of the indicated groups. n = 5 mice/group. **E,** (Upper panel) Representative WGA-stained RV sections to assess hypertrophy of cardiac myocytes. Scale bar: 50 µm. (Lower panel) Quantitative analysis. n = 5 mice/group. **F,** (Upper panel) Representative H&E-stained pulmonary artery sections from the indicated groups. (Lower panel) Percentage of wall thickness of small arteries in relation to cross-sectional diameter. n = 5 mice/group. **G,** Design of the study. Ten-week-old C57B6 mice subcutaneously received 20 mg/kg of SU5416 (SU) and were then exposed to two weeks of chronic hypoxia. SU5416 was injected once a week during the next two weeks. Mice were then randomly assigned to receive every 3 days a dose of recombinant BMP3 (0.3 mg/kg) or a saline solution (as control) at week 12 for 2 weeks. The endpoint for hemodynamic measurements and sacrifice was at week 14. **H,** RVSP of the indicated groups. n = 5-6 mice/group. I, Right ventricular weight of the indicated groups. n = 5-6 mice/group. **J,** Fulton index of the indicated groups. n = 5-6 mice/group. **K,** (Upper panel) Representative WGA-stained RV sections to assess hypertrophy of cardiac myocytes. Scale bar: 50 µm. (Lower panel) Quantitative analysis. n = 5-6 mice/group. **L,** (Upper panel) Representative H&E-stained pulmonary artery sections from the indicated groups. (Lower panel) Percentage of wall thickness of small arteries in relation to cross-sectional diameter. n = 5-6 mice/group. * P < 0.05; ** P < 0.01; *** P < 0.001, by 1-way ANOVA

**Figure S5:**
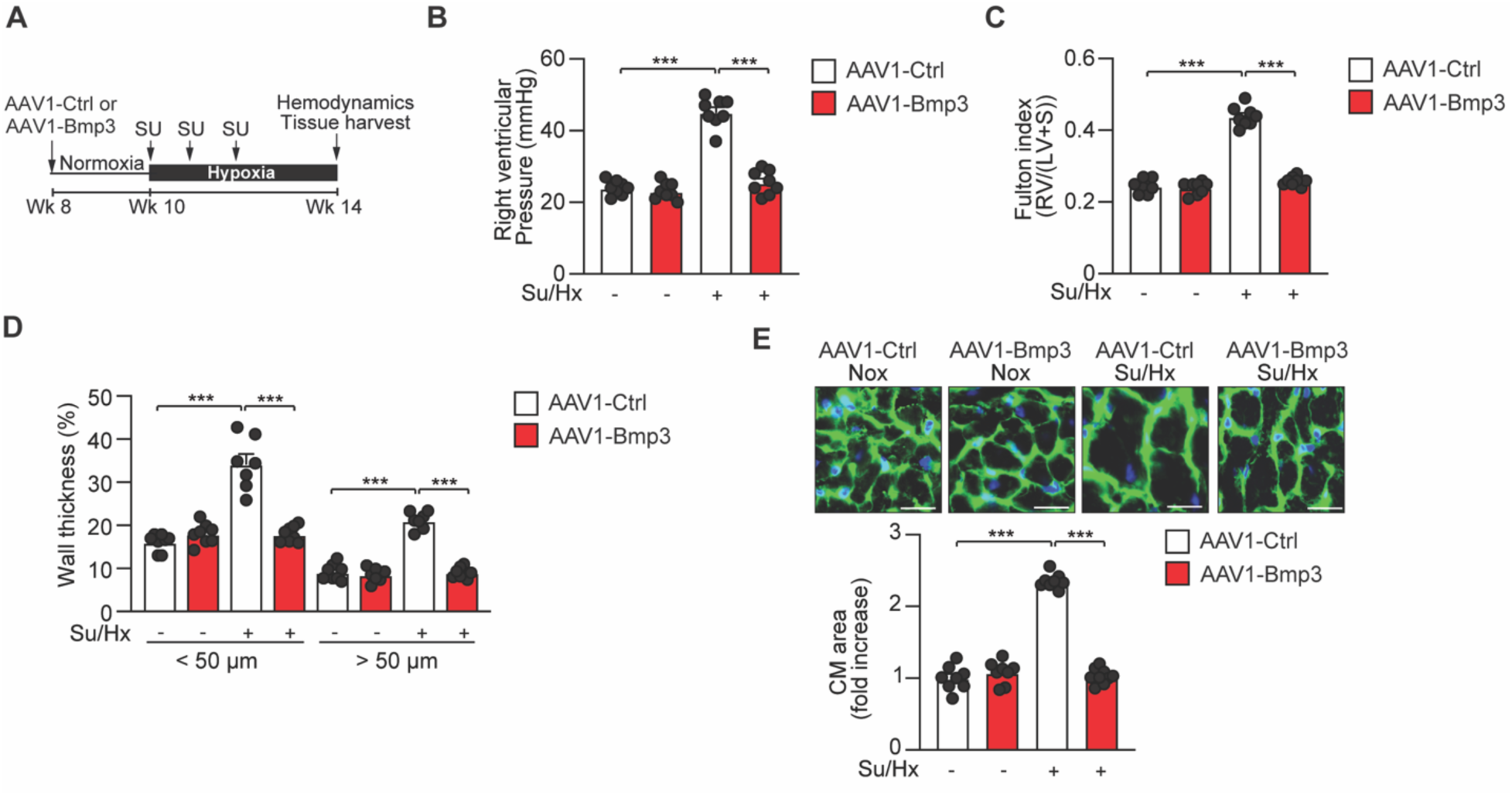
A lung specific overexpression of BMP3 prevents PH in female mice. **A,** Design of the study. Female mice were randomly assigned to receive AAV1-Ctrl or AAV1-Bmp3 at week 8. Two weeks later, mice received 20 mg/kg of SU5416, and were then exposed to four weeks of chronic hypoxia. SU5416 was injected once a week during the next two weeks. The endpoint for hemodynamic measurements and sacrifice was at week 14. **B,** RVSP of the indicated groups. n = 6-8 mice/group. **C,** Fulton index of the indicated groups. n = 6-8 mice/group. **D,** Percentage of arterial wall thickness in relation to cross-sectional diameter. n = 6-8 mice/group. **E,** (Upper panel) Representative WGA-staining of RV sections. Scale bar: 25 µm. (Lower panel) Quantitative analysis. Scale bar: 25 µm. n = 6-8 mice/group. *** P < 0.001, by 2-way ANOVA. Su/Hx: Sugen/Hypoxia, CM: Cardiomyocyte.

**Figure S6:**
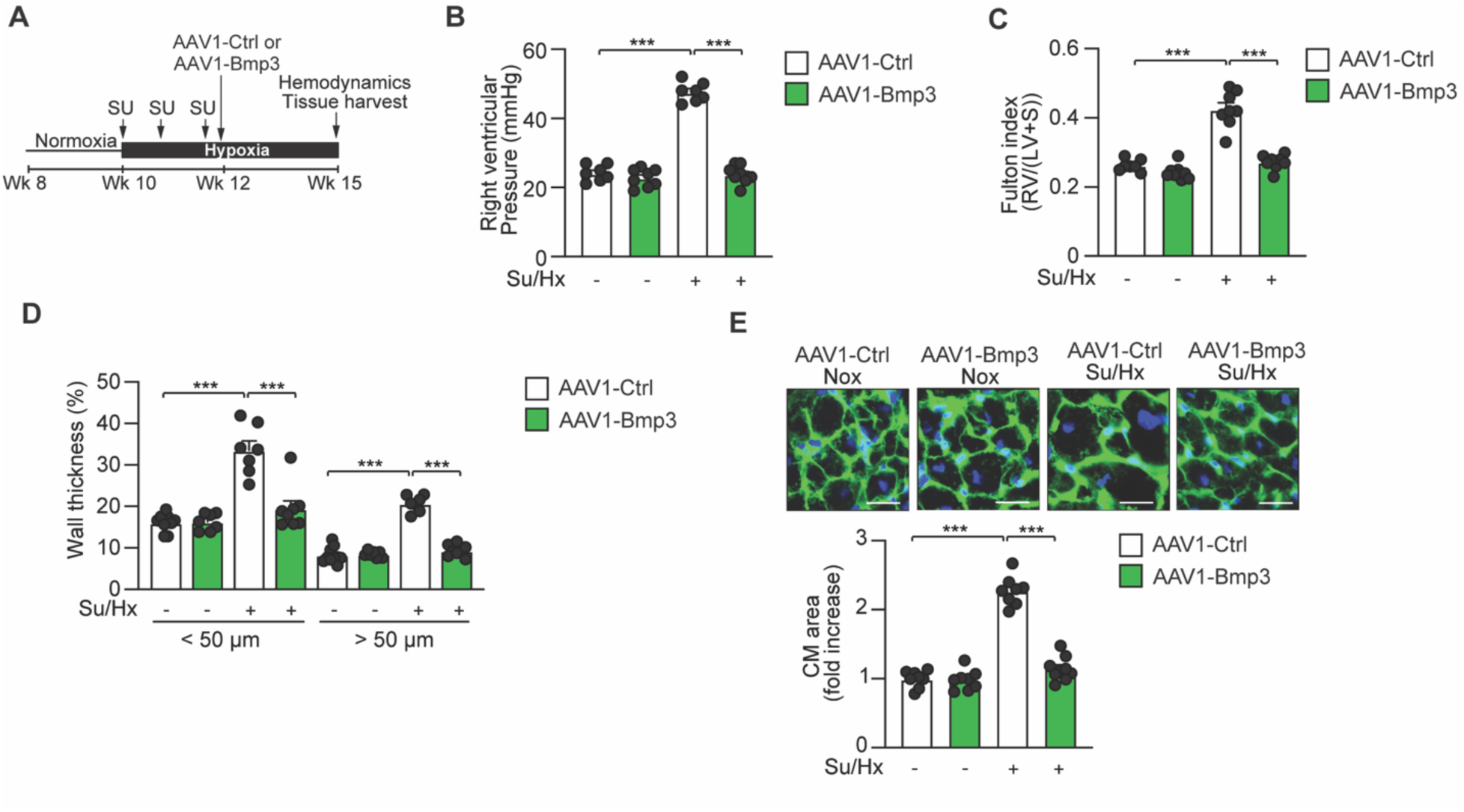
A lung specific overexpression of BMP3 reverses PH in female mice. **A,** Design of the study. Female mice received 20 mg/kg of SU5416, and were then exposed to two weeks of chronic hypoxia. SU5416 was injected once a week during the next two weeks. Mice were then randomly assigned to receive AAV1-Ctrl or AAV1-Bmp3 at week 12. The end point for hemodynamic measurements and sacrifice was at week 16. **B,** RVSP of the indicated groups. n= 5-8 mice/group. **C,** Fulton index of the indicated groups. n= 5-8 mice/group. **D,** Percentage of arteries wall thickness in relation to cross-sectional diameter. n= 5-8 mice/group. **E,** (Upper panel) Representative WGA-staining of RV sections. Scale bar: 25 µm. (Lower panel) Quantitative analysis. n= 5-8 mice/group. *** P < 0.001, by 2-way ANOVA. Su/Hx: Sugen/Hy­ poxia, CM: Cardiomyocyte.

## SUPPLEMENTAL TABLES

**Table S1.**
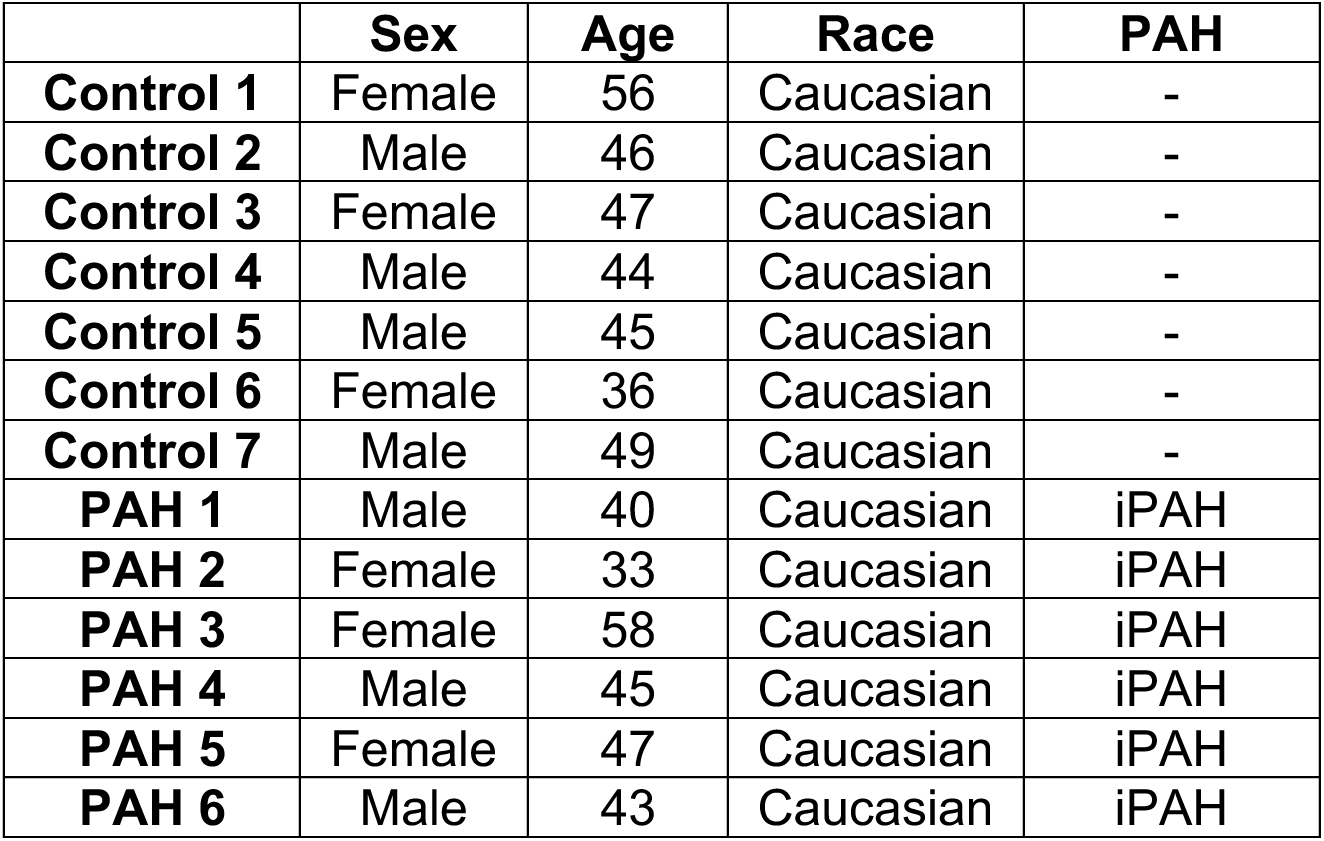
Detailed PASMC characteristics used in this study.

|  | **Sex** | **Age** | **Race** | **PAH** |
| --- | --- | --- | --- | --- |
| **Control 1** | Female | 56 | Caucasian | - |
| **Control 2** | Male | 46 | Caucasian | - |
| **Control 3** | Female | 47 | Caucasian | - |
| **Control 4** | Male | 44 | Caucasian | - |
| **Control 5** | Male | 45 | Caucasian | - |
| **Control 6** | Female | 36 | Caucasian | - |
| **Control 7** | Male | 49 | Caucasian | - |
| **PAH 1** | Male | 40 | Caucasian | iPAH |
| **PAH 2** | Female | 33 | Caucasian | iPAH |
| **PAH 3** | Female | 58 | Caucasian | iPAH |
| **PAH 4** | Male | 45 | Caucasian | iPAH |
| **PAH 5** | Female | 47 | Caucasian | iPAH |
| **PAH 6** | Male | 43 | Caucasian | iPAH |

**Table S2.**
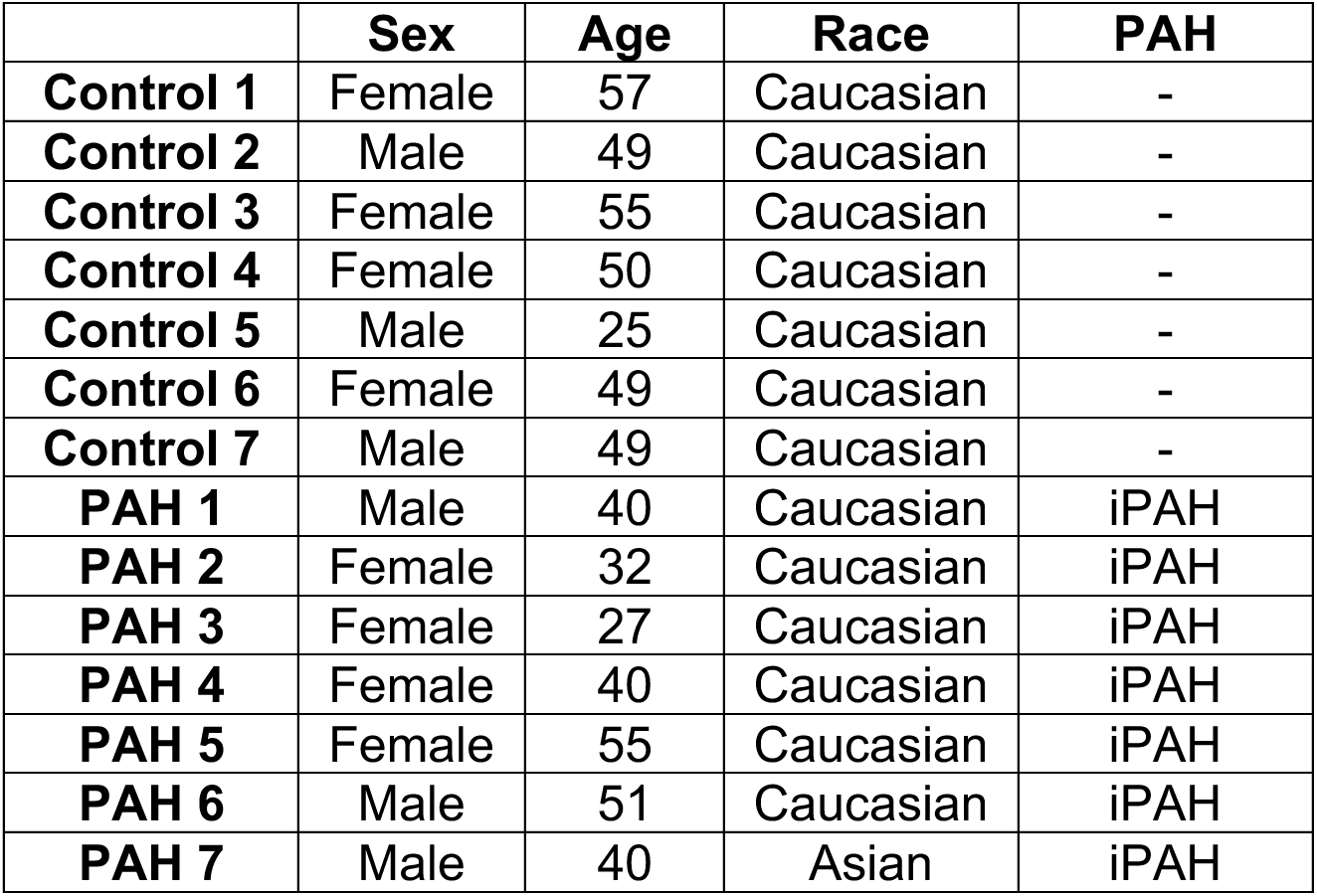
Detailed PAEC characteristics used in this study.

|  | **Sex** | **Age** | **Race** | **PAH** |
| --- | --- | --- | --- | --- |
| **Control 1** | Female | 57 | Caucasian | - |
| **Control 2** | Male | 49 | Caucasian | - |
| **Control 3** | Female | 55 | Caucasian | - |
| **Control 4** | Female | 50 | Caucasian | - |
| **Control 5** | Male | 25 | Caucasian | - |
| **Control 6** | Female | 49 | Caucasian | - |
| **Control 7** | Male | 49 | Caucasian | - |
| **PAH 1** | Male | 40 | Caucasian | iPAH |
| **PAH 2** | Female | 32 | Caucasian | iPAH |
| **PAH 3** | Female | 27 | Caucasian | iPAH |
| **PAH 4** | Female | 40 | Caucasian | iPAH |
| **PAH 5** | Female | 55 | Caucasian | iPAH |
| **PAH 6** | Male | 51 | Caucasian | iPAH |
| **PAH 7** | Male | 40 | Asian | iPAH |

**Table S3.** Hemodynamic data of the sham and PH diseased pigs.

|  | **Mean PAP**<br>**(mmHg)** | **RVSP**<br>**(mmHg)** | **CO**<br>**(l/min)** |
| --- | --- | --- | --- |
| Pig Sham 1 | 17 | 29 | 2.7 |
| Pig Sham 2 | 17 | 43 | 2.1 |
| Pig Sham 3 | 14 | 30 | 2.5 |
| Pig Sham 4 | 10 | 28 | 2.4 |
| Pig Sham 5 | 18 | 34 | 2.8 |
| Pig Sham 6 | 21 | 32 | 4.1 |
| Pig PH 1 | 68 | 90 | 3.3 |
| Pig PH 2 | 47 | 58 | 3.2 |
| Pig PH 3 | 52 | 75 | 5.1 |
| Pig PH 4 | 60 | 88 | 4.9 |

Mean PAP: Mean pulmonary arterial pressure, RVSP: Right ventricular systolic pressure, CO: Cardiac output.

**Table S4.**
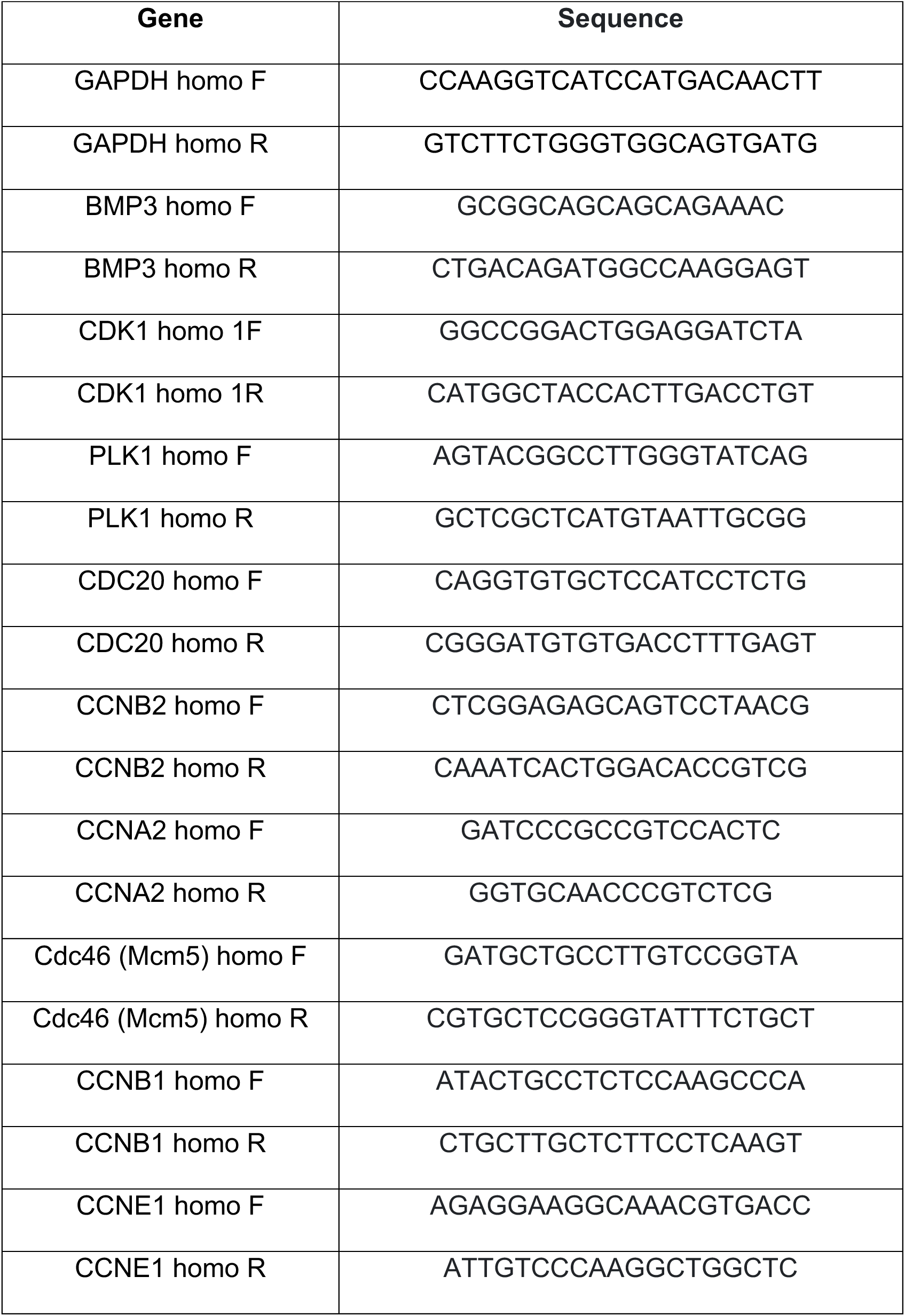
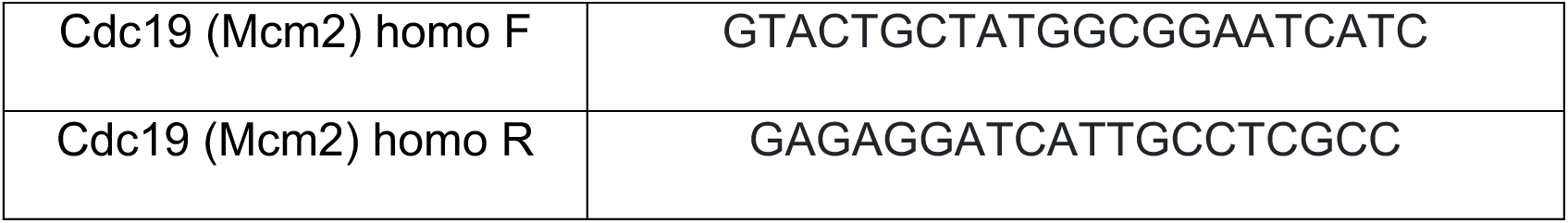
Primer sequences used for human gene expression analysis.

**Table S5.**
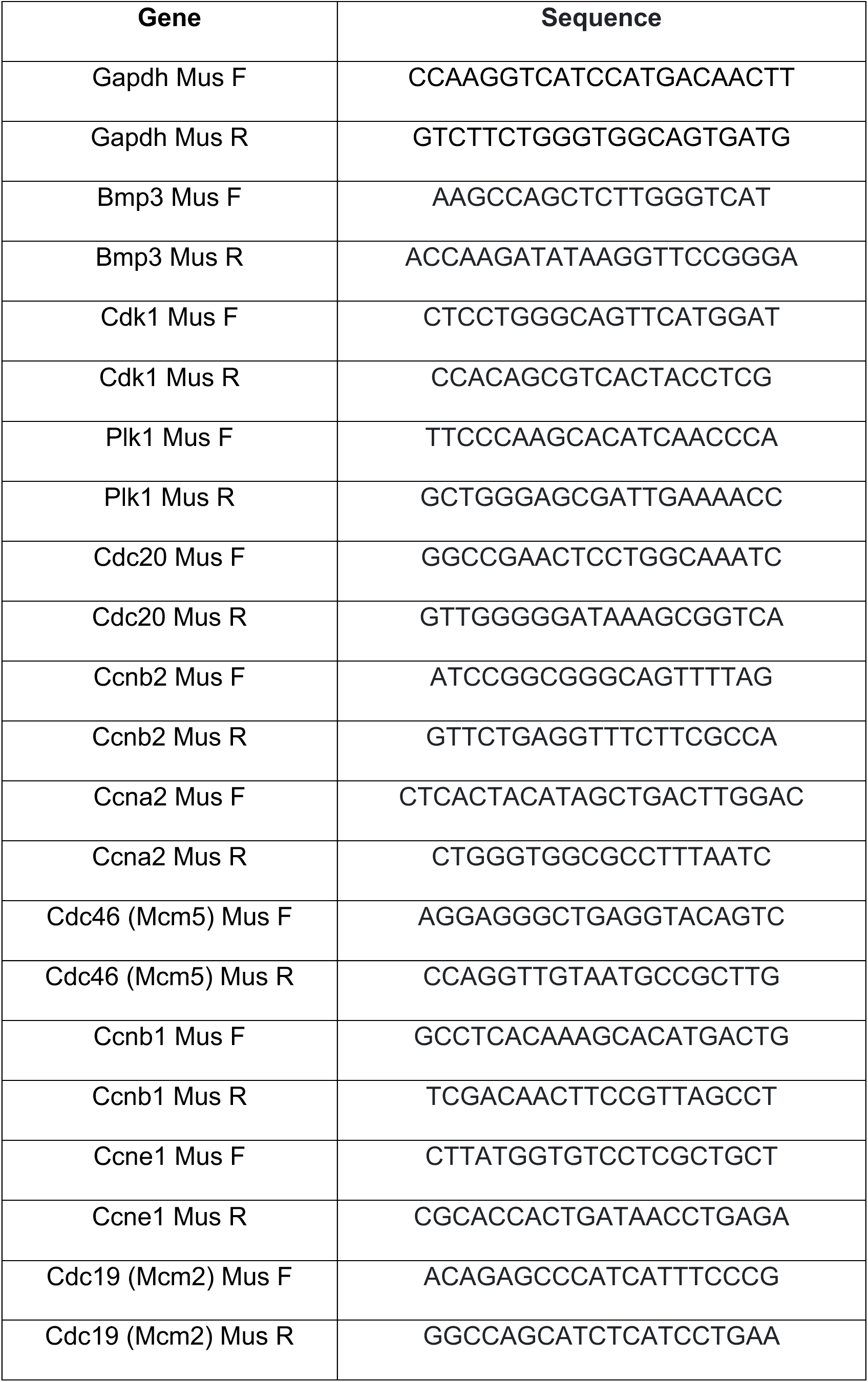
Primer sequences used for mouse gene expression analysis.

